# MERS-CoV antagonizes PKR activation by inhibiting its condensation at viral replication complexes

**DOI:** 10.1101/2025.08.18.670656

**Authors:** Ebba K. Blomqvist, Nicole Bracci, Helena Winstone, J. Monty Watkins, Susan R. Weiss, James M. Burke

## Abstract

Middle East respiratory syndrome coronavirus (MERS-CoV) is a highly pathogenic virus that antagonizes innate immune responses, including the protein kinase R (PKR) pathway. Here, we examine the process of PKR activation in response to an immunostimulatory MERS-CoV mutant encoding an inactive endoribonuclease U and a deletion of accessory protein NS4a. We show that PKR condenses and activates on viral dsRNA proximal to viral double-membrane vesicles (DMVs). Condensates composed of activated PKR disassociate from dsRNA and dissolve, releasing activated PKR molecules into the cytosol where they phosphorylate eIF2α to initiate the integrated stress response. MERS-CoV NS4a protein prevents PKR activation by condensing on dsRNA and occluding PKR binding. Lastly, PKR condensation coincided with its activation in response to Zika virus. These findings establish a comprehensive model for PKR activation in response to positive-strand RNA viruses that replicate within membrane-associated complexes and elucidate how MERS-CoV antagonizes this crucial antiviral pathway.

**GRAPHICAL ABSTRACT:** 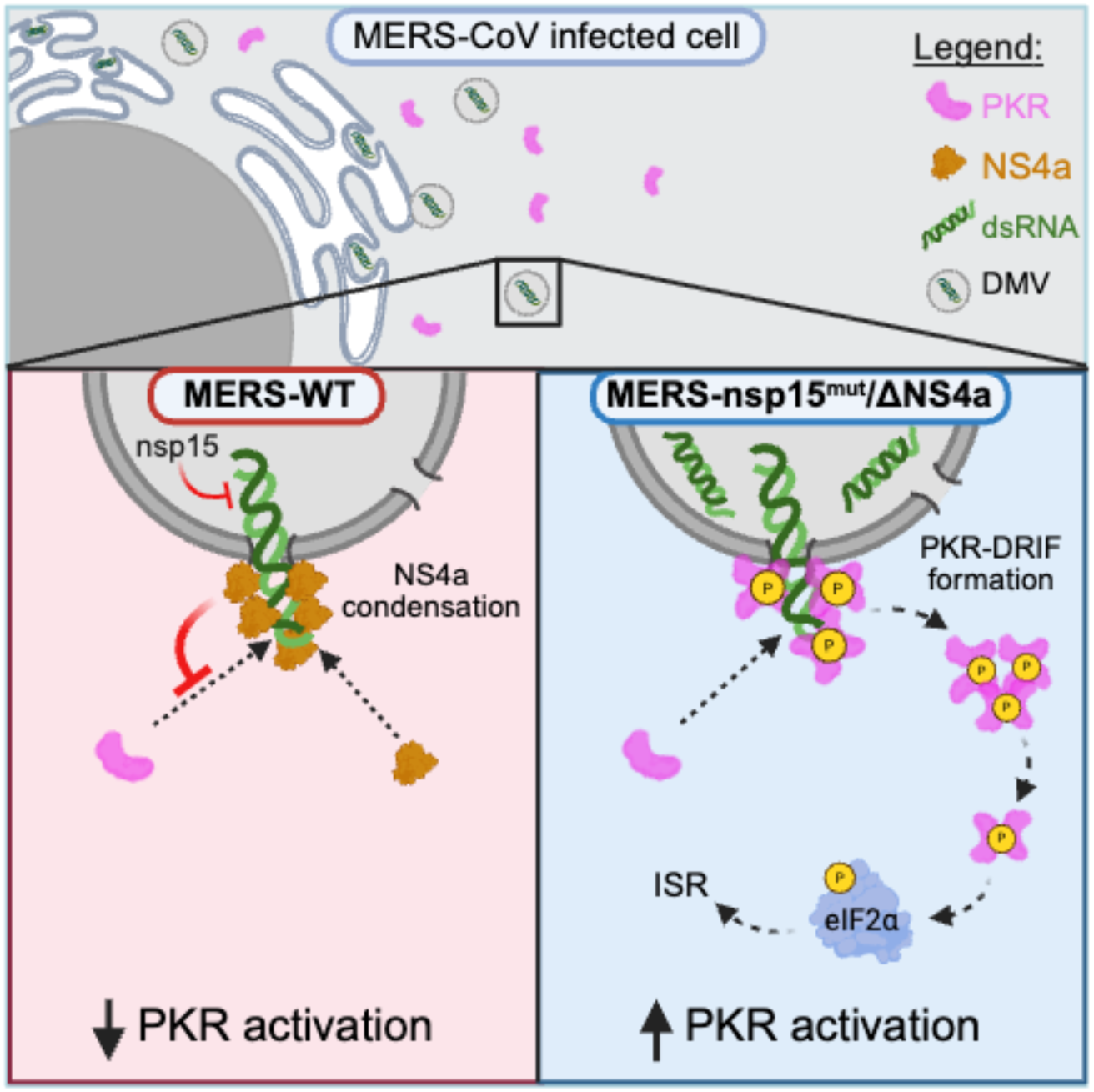

**HIGHLIGHTS:**

- PKR condenses on viral dsRNA exposed at membrane-associated replication complexes
- PKR condensation initiates autophosphorylation of PKR
- Activated p-PKR molecules disassociate from dsRNA and re-localize to the cytosol
- MERS-CoV NS4a inhibits PKR activation by occluding PKR condensation on dsRNA.

## INTRODUCTION

Middle East respiratory syndrome coronavirus (MERS-CoV) has a case fatality rate of 35% according to the World Health Organization (WHO), and is considered a pathogen with pandemic potential^1,2^. A contributing factor proposed for the high virulence of MERS-CoV is that it inhibits cell autonomous innate immune responses^3–5^, thus enhancing viral replication and dissemination. MERS-CoV inhibits innate immune responses via two general mechanisms. First, it replicates its RNA within double membrane vesicles (DMVs) derived from rearranged endoplasmic reticulum membranes^6^. This confines dsRNA generated during RNA replication within membranous structures to prevent cytosolic pattern recognition receptors (PRRs) from binding to dsRNA and initiating antiviral signaling cascades. Second, MERS-CoV encodes multiple proteins, including nsp15 and NS4a, known to inhibit innate immune responses^7,8^. Nsp15 contains an endoribonuclease (EndoU) conserved among coronaviruses that limits dsRNA accumulation by cleaving viral ssRNA or bulged dsRNA^9–13^. NS4a encoded by MERS-CoV and related merbecoviruses is thought to limit PRR activation via its dsRNA-binding domain^7,14^, though the precise mechanism by which NS4a limits PRR activation has not formally been demonstrated.

Protein kinase R (PKR) is a PRR that antagonizes the replication of numerous viruses. PKR binds to viral dsRNA via its N-terminal domain^15,16^. Concentration of PKR on dsRNA promotes its homodimerization, which leads to autophosphorylation (p-PKR) and activation of its C-terminal kinase domain^16–18,19^. Activated PKR (p-PKR) phosphorylates proteins that initiate stress and inflammatory responses ^20–22^. A primary target of PKR is the translation initiation factor, eIF2α. PKR phosphorylates eIF2α on serine 51 (p-eIF2α)^23^. This activates the integrated stress response (ISR), which arrests canonical translation initiation of most host and viral mRNAs and triggers the assembly of stress granules^24,25^. Consequently, many viruses such as MERS-CoV inhibit the activation of PKR^7,26^.

Despite greater than four decades of research on PKR, fundamental aspects of PKR activation remain unresolved, including where PKR interacts with viral dsRNA (in the cytosol or within DMVs), whether activated PKR remains associated with dsRNA or disassociates, and how viral proteins antagonize PKR activation. An important advancement in understanding PKR activation was the observation that PKR and dsRNA assemble into condensates termed dsRNA-induced foci (DRIF)^27^. PKR-DRIF have been observed during dengue virus serotype II (DENV-2) infection^28^, as well as measles virus^29^. It is unknown if PKR condensation on dsRNA is important for its activation during viral infection and whether viruses antagonize this process.

In this article, we investigate the subcellular location and molecular process of PKR activation in response to an immunostimulatory MERS-CoV mutant virus encoding loss of function mutations in EndoU and NS4a (MERS-nsp15^mut^/ΔNS4a)^7^. Using super-resolution microscopy, proximity ligation assays, and immunogold transmission electron microscopy, we show that PKR is activated in the majority of lung derived A549 cells infected with MERS-nsp15^mut^/ΔNS4a. The activation of PKR initiates upon its condensation on dsRNA immediately proximal to viral DMVs. Subsequently, PKR condensates disassociate from viral dsRNA/DMVs and dissolve, releasing activated p-PKR molecules into the cytosol where they phosphorylate eIF2α. MERS-CoV NS4a directly interferes with PKR activation by condensing on dsRNA, thus occluding PKR binding to dsRNA. Lastly, we show that PKR condensation coincides with its activation in response to Zika virus. These findings define the subcellular localization and molecular processes of PKR activation in response to positive-strand RNA viruses that replicate within membrane-associated replication complexes, such as coronaviruses and flaviviruses, and reveal the molecular mechanisms by which MERS-CoV antagonizes PKR activation.

## RESULTS

### MERS-CoV nsp15 and NS4a inhibit PKR-mediated translation arrest

To resolve the subcellular location and molecular processes of PKR activation in response to coronaviruses, as well as how MERS-CoV nsp15 and NS4a antagonizes it, we developed an immunofluorescence assay (IFA) for p-PKR to visualize activated PKR by microcopy. We infected human alveolar epithelial A549 cells expressing dipeptidyl peptidase-4 (DPP4) (hereby referred to as A549) with an immunostimulatory MERS-CoV mutant virus that encodes an enzymatically inactive nsp15 and deletion of NS4a (MERS-nsp15^mut^/ΔNS4a), which has previously been shown to trigger PKR activation^7,26^. We co-stained cells for dsRNA (infection marker).

As expected, wild-type MERS-CoV (MERS-WT) did not frequently induce PKR activation, as only a small percentage (<5%) of infected (dsRNA-positive) cells stained for p-PKR above levels observed in mock-infected cells (Figures 1A and S1A). In contrast, MERS-nsp15^mut^/ΔNS4a induced frequent and robust PKR activation in most infected cells (83%) (Figures 1A and S1B). We confirmed that the p-PKR staining was specific to PKR, as it was not observed in PKR-KO cells (Figure S1C), and western blot analyses confirmed the increase in p-PKR in response to MERS-nsp15^mut^/ΔNS4a (Figure S1D). Both MERS-nsp15^mut^ and MERS-ΔNS4a induced more frequent and robust PKR activation than MERS-WT, though neither triggered PKR activation as frequently or robustly as MERS-nsp15^mut^/ΔNS4a (Figure 1A).

**Figure 1.**
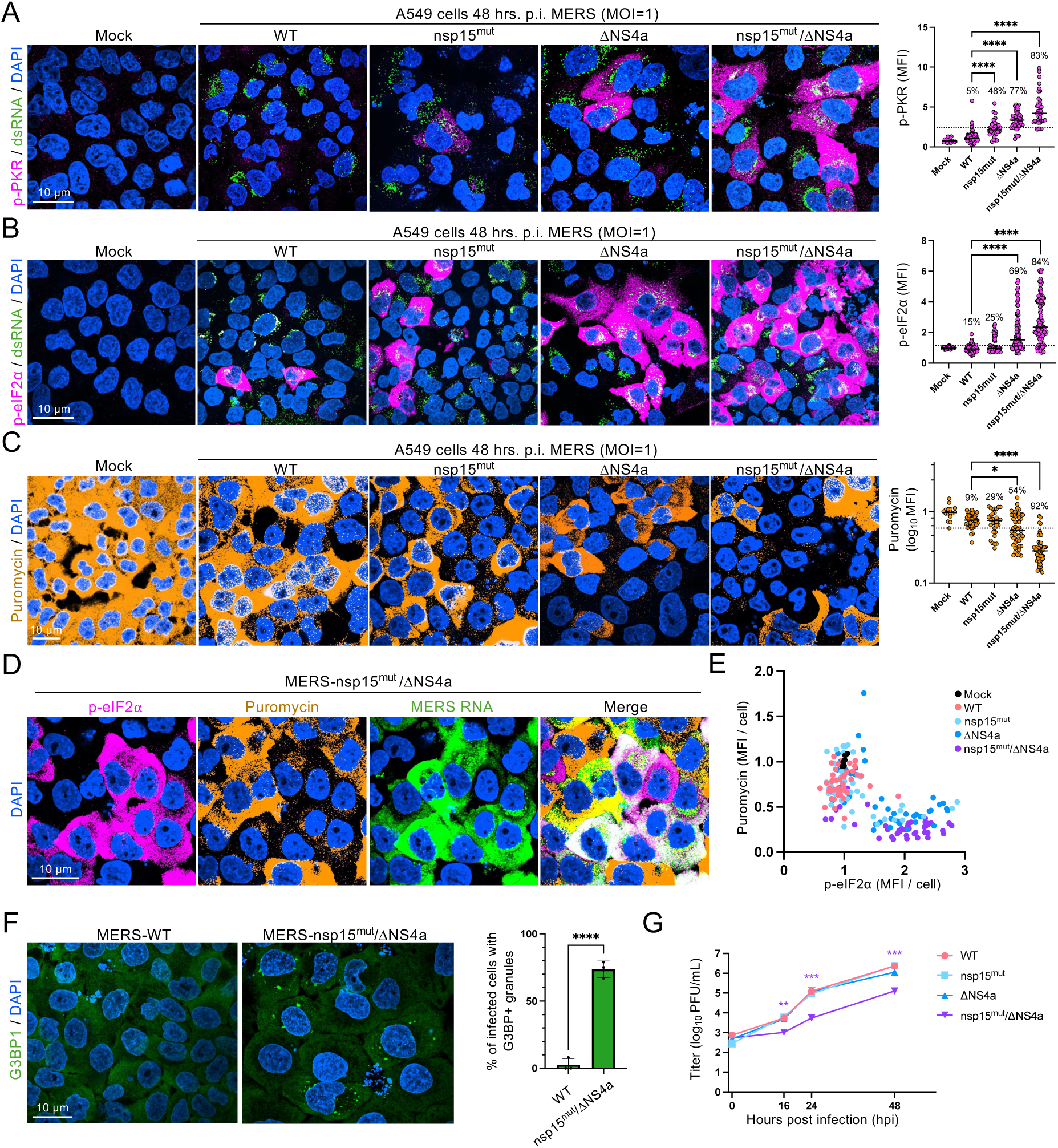
MERS-CoV nsp15 and NS4a inhibit PKR-mediated arrest of translation. (A) Immunofluorescence assay (IFA) for p-PKR and dsRNA in A549-DPP4 cells 48 hours post-infection with the indicated MERS-CoV strains. The graph on the right is the quantification of cytosolic p-PKR intensity per cell (dot). The dotted line indicates the cutoff (two-fold over average intensity in mock cells) for determining if a cell as has activated PKR. (B) IFA for phosphorylated eIF2⍺ (p-eIF2⍺) and dsRNA. (C) RiboPuromycylation immunofluorescence assay. Quantification of cytosolic puromycin intensity per cell (dot) as represented in images depicting the percentage of cells with puromycin below mock (dotted line). (D) IFA for eIF2⍺ (magenta) and puromycin (orange) combined with smFISH probing for 5’-MERS RNA. (E) Scatter plot showing p-eIF2⍺ levels compared to puromycin levels in individual cells (dot) infected with the indicated MERS virus for 48 hours. (F) IFA for G3BP1 (stress granule marker) 48 hours p.i. with either WT MERS or MERS-nsp15^mut^/ΔNS4a. The graph shows the percentage of infected cells containing G3BP1-granules from three fields of view. (G) Quantification of viral production at indicated times post-infection by plaque assays.

The enhanced activation of PKR in response to MERS-nsp15^mut^/ΔNS4a was sufficient to trigger the integrated stress response (ISR) in most infected cells based on the following observations. First, cells infected with MERS-nsp15^mut^/ΔNS4a displayed robust phosphorylation of eIF2α (Figure 1B), which was PKR-dependent as it was abolished in PKR-KO cells (Figure S1E). Second, RiboPuromycylation assays showed that cells infected with MERS-nsp15^mut^/ΔNS4a displayed a reduction in bulk translation in comparison to cells infected with MERS-WT (Figure 1C). Notably, the frequency of both phosphorylation of eIF2α and translation repression was proportional to the frequency of p-PKR observed in each respective viral strain (Figures 1A-C), and the reduction in bulk translation correlated with p-eIF2α abundance (Figures 1D-E). Lastly, most cells infected with MERS-nsp15^mut^/ΔNS4a assembled stress granules, which did not assemble in response to MERS-WT (Figures 1F and S2A).

Quantification of viral replication showed that MERS-nsp15^mut^/ΔNS4a displayed a significant (10-fold) reduction in viral replication in comparison to MERS-WT and the single mutants (Figure 1G). Combined, these data indicate that nsp15 and NS4a synergistically antagonize the activation process of PKR, resulting in the inhibition of the ISR and the enhancement of viral replication capacity, consistent with previous studies ^7,26^. Importantly, these data demonstrate that we can visualize PKR activation during MERS-nsp15^mut^/ΔNS4a infection.

### PKR activates upon condensation at sites of MERS-nsp15^mut^/ΔNS4a replication complexes

We next examined the subcellular location at which PKR binds and activates on dsRNA, which we reasoned could occur on dsRNA associated with viral double membrane vesicle (DMV) containing replication-transcription complexes (RTC) or on dsRNA that leaks out of DMVs and diffuses into the cytosol.

A key observation we made is that activated PKR (p-PKR), in addition to being diffuse throughout the cytosol, aggregated in puncta that were readily observed with low intensity projection (Figure 2A). Despite reports that PKR localizes to stress granules ^30,31^, IFA for G3BP1 (stress granule and RNase L-induced body marker) showed that the p-PKR puncta did not co-localize with either stress granules (Figure S2B) or RNase L-induced bodies (RLB), which are stress granule-like condensates that form upon RNase L activation and contain G3BP1 (Figure S2C).

**Figure 2.**
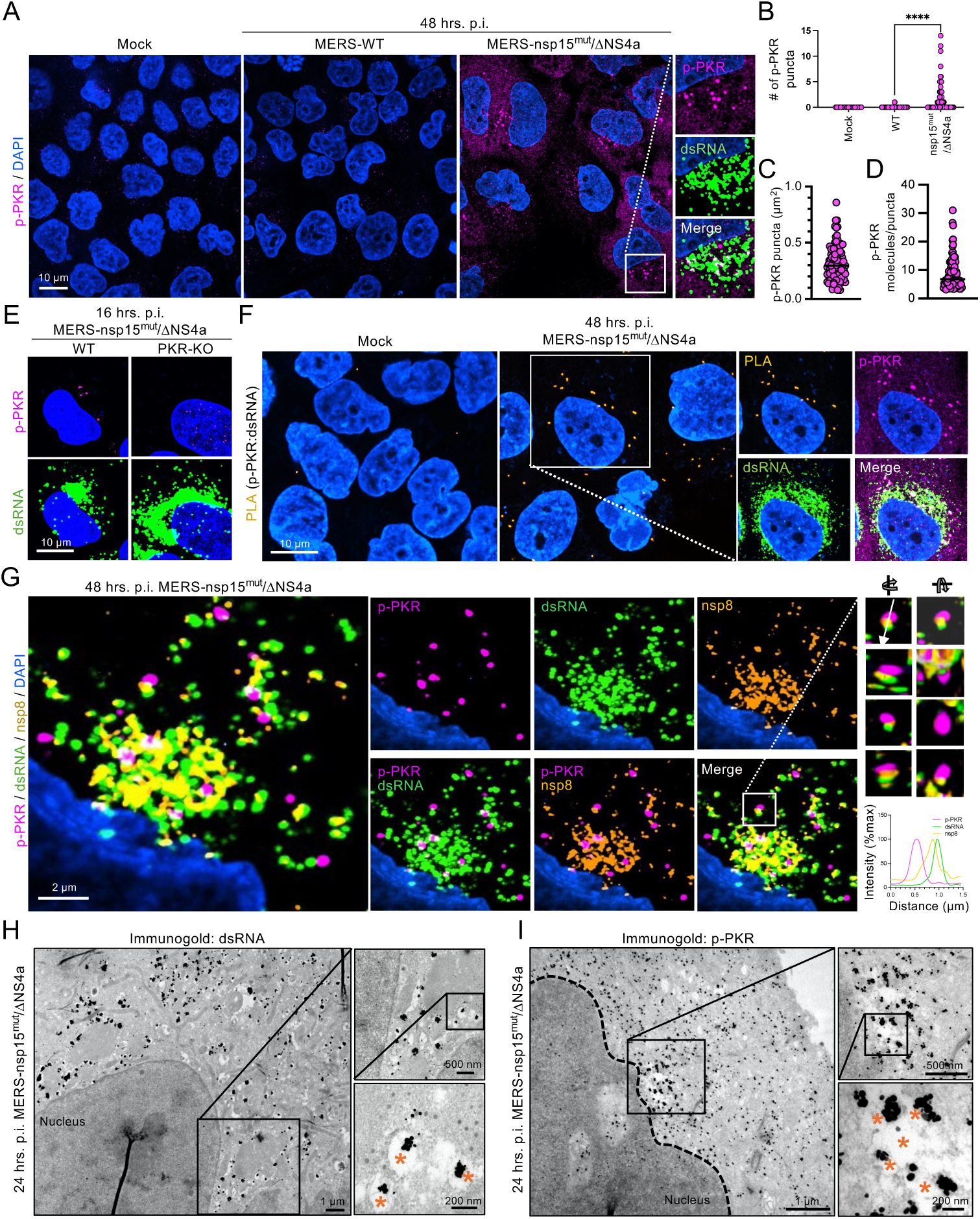
PKR activates upon condensation at viral DMV replication complexes. (A) IFA for p-PKR and dsRNA 48 hours p.i. The large image panels only display p-PKR staining at low intensity projections to present the concentration of p-PKR in puncta that co-localize with dsRNA (shown in the inset). (B) Quantification of the number of PKR puncta per cell (dot). (C) Quantification of area of PKR puncta in MERS-nsp15^mut^/ΔNS4a infected cells (each dot represents one PKR puncta). (D) Estimated number of p-PKR dimers contained in each PKR puncta in MERS-nsp15^mut^/ΔNS4a infected cells (each dot represents one PKR puncta). (E) IFA for p-PKR and dsRNA 16 hours p.i. A549-DPP4 and A549-DPP4-PKR KO cells infected with MERS-nsp15^mut^/ΔNS4a. (F) Co-IFA and PLA for p-PKR (magenta) and dsRNA (green) in A549-DPP4 infected cells. The large image panels only p-PKR : dsRNA PLA signal, whereas the inset shows the relative location of PLA signal with p-PKR and dsRNA. (G) Super-resolution microscopy of IFA for p-PKR, dsRNA, and nsp8 48 hours p.i. with MERS-nsp15^mut^/ΔNS4a infected A549-DPP4 cells. The intensity threshold was set to specifically visualize the p-PKR puncta. Image panels showing individual and dual IFA stains are to the right of the large composite image. 3-D visualization on the inset was performed, as well as an intensity profile along the line, which is graphed in the lower right. (H-I) Immunogold TEM of A549-DPP4 cells at 24 p.i. DMVs are indicated with orange asterisks. (H) dsRNA staining (I) p-PKR staining.

The assembly of the p-PKR puncta coincides with PKR activation, as they were not observed in mock-infected cells (Figure 2B). The p-PKR puncta were mostly localized near the perinuclear region of the cell where viral dsRNA concentrated (Figure 2A). Because this is presumably where individual viral DMV-RTCs concentrate on the ER, we refer to this region generally as the viral region of replication (ROR). The p-PKR puncta are specific to PKR, as they were not observed in PKR-KO cells infected with MERS-nsp15^mut^/ΔNS4a (Figure S3A), and we observed them by IFA for total PKR (Figure S3B). The average number of p-PKR puncta per MERS-nsp15^mut^/ΔNS4a-infected cell was variable, between 2-10 puncta (Figure 2B). All p-PKR puncta were spherical and similar in size (0.3 µm^2^) (Figure 2C). By calculating p-PKR intensity in the cytosol versus the p-PKR puncta, we estimate that each p-PKR punctum has between 3-30 p-PKR molecules (Figure 2D).

We hypothesized that the p-PKR puncta were involved in the activation process of PKR. Three key observations support that the p-PKR puncta are the sites of PKR activation on dsRNA. First, IFA for p-PKR at 16 hours p.i. showed that p-PKR primarily localized to punctate structures within the ROR as opposed to the cytosol (Figure 2E), indicating that these are the initial sites of PKR activation. Second, proximity-ligation assay (PLA) between dsRNA and PKR at 16 hours p.i. showed that dsRNA : PKR interactions were typically localized to the ROR (Figure S4A). Third, we performed PLA for dsRNA and p-PKR combined with IFA at 48 hours p.i., which is when p-PKR is localized through the cytosol as well as in puncta. These analyses showed that interactions between p-PKR and dsRNA occurred at p-PKR puncta co-localized with dsRNA in the ROR, whereas p-PKR diffusely localized in the cytosol was not associated with detectable dsRNA interaction by PLA (Figure 2F). Notably, mock-infected cells displayed minimal PLA (dsRNA : p-PKR) puncta, suggesting our PLA signal is dependent on dsRNA (Figure 2F). Moreover, the proximity ligation was PKR-dependent, as it was abolished in PKR-KO cells (Figure S4B). Combined, these data indicate that the p-PKR puncta are the sites at which PKR binds and activates on viral dsRNA.

We next asked if the p-PKR puncta are associated with dsRNA that had leaked out into the cytoplasm or whether the p-PKR puncta were associated with DMVs in which viral RNA replication occurs. To do this, we performed super-resolution fluorescence microscopy to determine if the p-PKR puncta co-localize with MERS-CoV nsp8, which specifically localizes to viral DMVs ^32,33^. Importantly, the p-PKR puncta typically co-localized with both dsRNA and nsp8 (Figure 2G). We note that some p-PKR puncta did not co-localize with either dsRNA or nsp8 (Figure 2G), which we address in the following section. Notably, nsp8 and dsRNA more closely co-localized with one another in comparison to the p-PKR puncta (Figure 2G), suggesting that the p-PKR puncta are likely immediately proximal to DMV-RTCs.

To further determine whether the p-PKR puncta are within or proximal to DMVs, we performed transmission electron microscopy (TEM) and immunogold staining for either dsRNA or p-PKR in cells 24 hours p.i. We observed dsRNA immunogold particles within structures consistent with the expected size and morphology of CoV DMVs (∼150-500 nm)^34,35^. These structures are predominantly localized to the perinuclear region of cells (Figure 2H), consistent with our IFA analyses of dsRNA and nsp8 (Figure 2G). We did not observe these DMV structures containing dsRNA in mock-infected cells (Figures S5A and S5B), indicating that they are specific to MERS-CoV infection.

Importantly, we observed that p-PKR formed aggregates on the outer surface of viral DMVs (Figures 2I and S5B). The p-PKR aggregates primarily localized to the perinuclear region of the cell concentrated with DMVs (Figures 2I and S5B), whereas p-PKR localized in the cytosol was composed of 1-2 immunogold particles. The immunogold signal for p-PKR was PKR dependent as it was abolished in A549-PKR-KO cells (Figure S5C). Combined, these data argue that PKR activates upon its condensation on dsRNA associated with viral DMVs. Because antiviral sensors condensing on dsRNA have been referred to as DRIF (<u>d</u>s<u>R</u>NA-induced foci) ^27–29,36^, we refer to the p-PKR puncta as PKR-DRIF.

### PKR-DRIF disassociate from dsRNA and subsequently disassemble

Because some PKR-DRIF were not associated with either dsRNA or nsp8 (Figure 2G), we considered the possibility that PKR-DRIF disassociate from dsRNA/DMVs. Three observations support this hypothesis. First, we verified that PKR-DRIF lack dsRNA by performing PLA for p-PKR and dsRNA. Indeed, the PKR-DRIF that did not co-localize with dsRNA lacked detectable interaction with dsRNA as assessed by PLA (dsRNA : p-PKR) (Figure 3A, inset 1). In contrast, PKR-DRIF that co-localized with dsRNA generated PLA (dsRNA : p-PKR) signal (Figure 3A, inset 2). These data support that some PKR-DRIF lack dsRNA. Second, we observed PKR-DRIF localized within the ROR that were not associated with DMVs (Figure 3B), suggesting that PKR-DRIF disassociate from DMVs. Importantly, we did not typically observe these dissociated PKR-DRIF localized outside the ROR where we observe p-PKR molecules (Figure 3B), suggesting that PKR-DRIF disassemble upon disassociation from DMVs.

**Figure 3.**
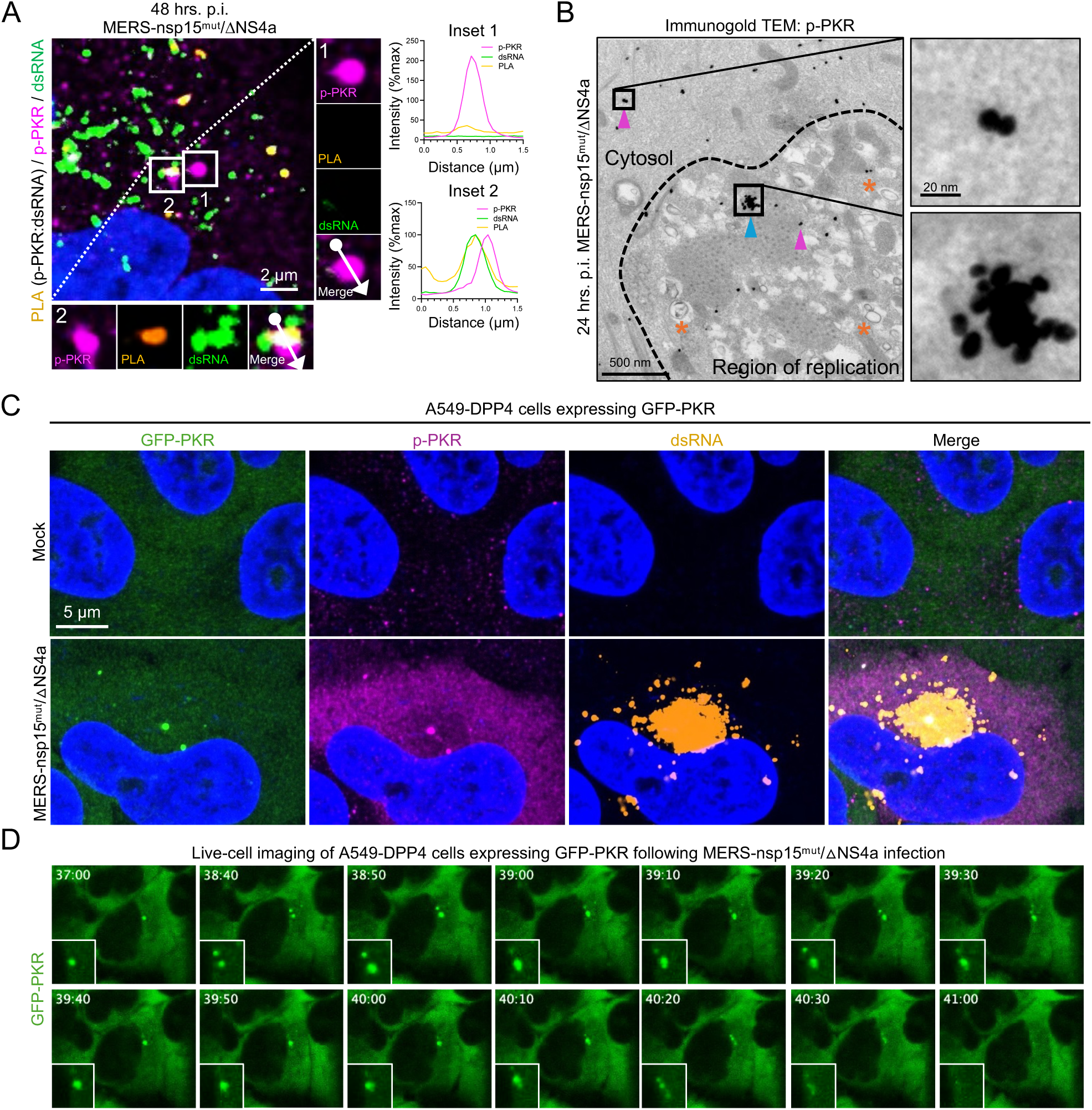
PKR-DRIPs dissociate from viral DMVs and dissolve, releasing activated PKR into the cytosol. (A) Co-IFA and PLA for p-PKR and dsRNA in A549-DPP4 cells at 48 hours p.i. with MERS-nsp15^mut^/ΔNS4a. Plot profile analyses were performed on PKR puncta in the indicated insets. (B) Immunogold TEM in A549-DPP4 cells stained for p-PKR. Pink arrows indicate p-PKR dimers, orange asterisks indicate viral DMVs, and the blue arrow indicates a PKR-DRIP. (C) IFA for dsRNA and p-PKR in A549-DPP4 cells expressing GFP-tagged PKR under mock conditions or 24 hrs. p.i. MERS-nsp15^mut^/ΔNS4a. (D) Live-cell imaging of A549-DPP4 cells expressing GFP-tagged PKR infected with MERS-nsp15^mut^/ΔNS4a (MOI 5) at the indicated time post-infection.

Third, we assessed whether PKR-DRIF undergo fission and disassembly in A549-DPP4 cell lines that stably express GFP-PKR, which formed puncta that co-stained for p-PKR within the ROR following MERS-nsp15^mut^/ΔNS4a infection (Figure 3C). Live cell imaging of following infection with MERS-nsp15^mut^/ΔNS4a showed that GFP-positive PKR-DRIF assemble between 12- and 24 hrs. p.i. in most cells infected with MERS-nsp15^mut^/ΔNS4a (Figure S6A and Supplemental Video S1). The GFP-PKR-DRIF could be observed through 48 hrs. p.i. (Figure S6B and Supplemental Video S2), indicating continuous assembly over the course of infection. Importantly, we observed that individual GFP-PKR-DRIF could undergo fission and dissolution over the course of viral infection (Figure 3D, Supplemental Video S3 and Figure S6C, Supplemental Video S4). These data demonstrate that PKR-DRIF disassemble and dissolve.

### PKR activates in the region of replication and re-localizes to the cytosol

Our above data support that PKR-DRIF disassociate from dsRNA/DMVs and dissolve in the ROR, releasing p-PKR molecules that then re-localize to the cytosol. Two observations further support this hypothesis. First, examination of the subcellular localization kinetics of p-PKR over the course of infection showed that p-PKR was primarily concentrated in the ROR at 16 hrs. p.i. (Figure 4A,B). At 24 hours p.i., we observed more diffuse p-PKR throughout the cytoplasm, which coincided with a reduction of p-PKR in the ROR (Figure 4A,B). By 48 hours p.i., most p-PKR was diffusely localized throughout the cytoplasm (Figure 4A,B). These data support that the activation of PKR initiates within the viral ROR and then re-localizes throughout the cytoplasm. Second, we analyzed the subcellular localization of p-PKR interactions by PLA. We observed an increase in PLA (p-PKR : eIF2α) in cells infected with MERS-nsp15^mut^/ΔNS4a in comparison to mock-infected cells (Figure 4C,D, S7A). Importantly, most PLA (p-PKR : eIF2α) signal was detected in the cytosol of the infected cells (Figure 4C,D and S7A,B). These data suggest that p-PKR primarily interacts with eIF2α following p-PKR re-localization from the ROR to the cytosol.

**Figure 4.**
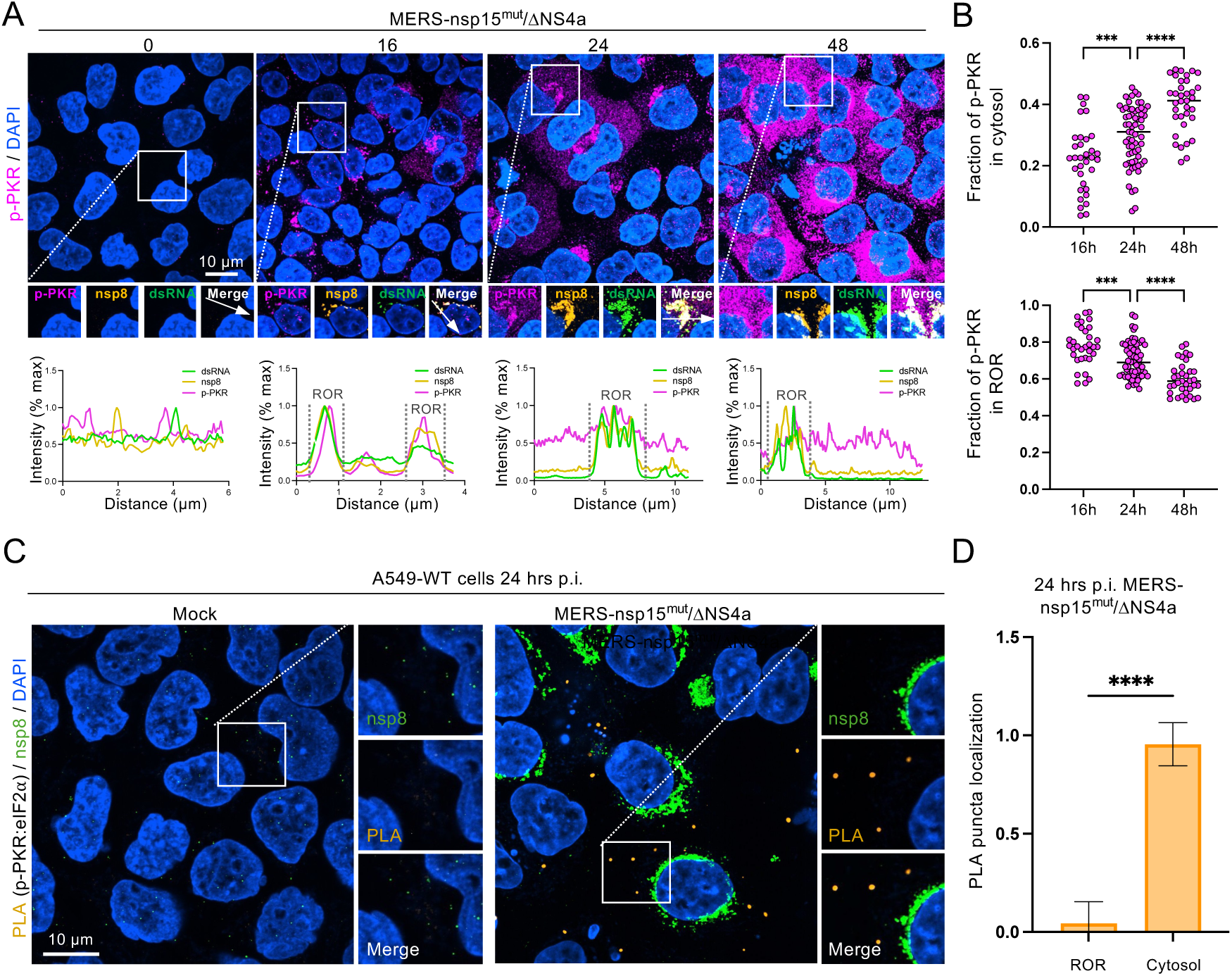
PKR activates in the region of replication and re-localizes to the cytosol. (A) IFA of A549-DPP4 cells either uninfected (0h) or infected with MERS-nsp15^mut^/ΔNS4a for 16, 24 and 48 hours. The cells are stained for p-PKR (magenta), nsp8 (orange) and dsRNA (green). Graphs at the bottom represent line traces from each time point, depicting the co-localization between each protein and dsRNA. (B) Quantification of panels from (A) where each dot represents one MERS-nsp15^mut^/ΔNS4a infected cell. (C) PLA between p-PKR and eIF2α 24 hours p.i. with MERS-nsp15^mut^/ΔNS4a in A549-WT cells. (D) Quantification of PLA puncta as represented in (C) analyzing three fields of view.

### NS4a condensation on dsRNA antagonizes PKR activation

We next asked why loss of nsp15 EndoU function and deletion of NS4a increased the activation of PKR. Previous studies have demonstrated that infection with catalytically inactivated EndoU during MERS-CoV ^7^ and SARS-CoV-2 ^8^ infections leads to higher levels of dsRNA compared to infections with WT-strains, and during murine hepatitis virus (MHV), infection with catalytically inactivated EndoU leads to increased cytosolic levels of dsRNA ^37^. Additionally, NS4a was shown to have a dsRNA-binding domain that antagonizes PRR activation ^7,14,26^. Therefore, we hypothesized that loss of nsp15 would lead to elevated dsRNA accumulation, and that a concurrent loss of NS4a would lead to an increase dsRNA-binding availability, which we could measure by IFA for dsRNA as shown in Figure 1A. Indeed, cells infected with MERS-WT displayed lower dsRNA intensity than those MERS-nsp15^mut^, MERS-ΔNS4a, or MERS-nsp15^mut^/ΔNS4a, and dsRNA IFA intensity was highest in cells infected with MERS-nsp15^mut^/ΔNS4a (Figure 5A). These data suggest that the nsp15 and NS4a limit dsRNA loads and/or reduce the availability of PKR to bind dsRNA.

**Figure 5.**
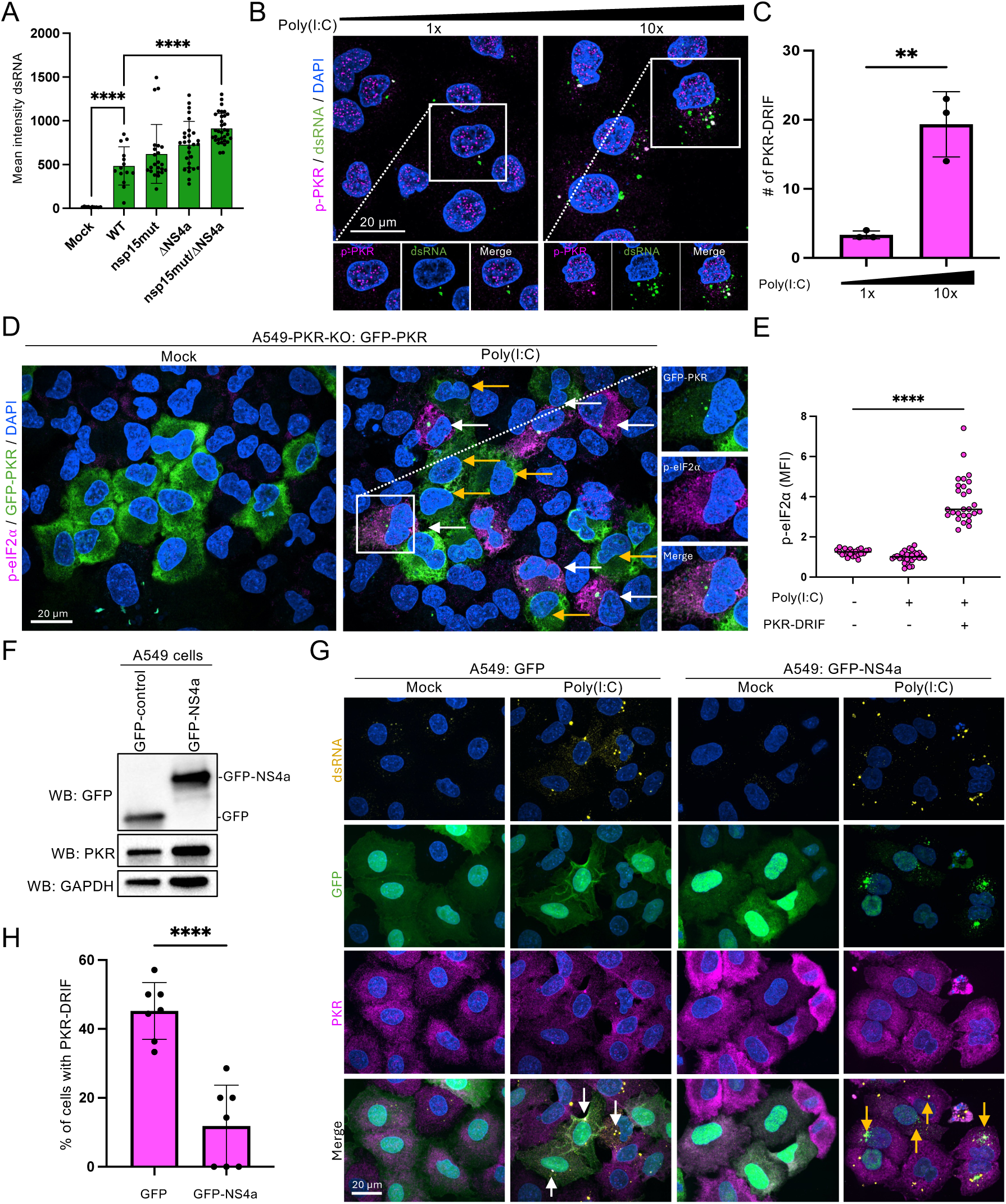
NS4a antagonizes PKR activation by binding dsRNA. (A) Quantification of dsRNA levels measured from IFA images of MERS-infected cells from Figure 1A. (B) IFA for p-PKR (magenta) and dsRNA (green) on cells treated with either 1x (250 ng) or 10x (2.5 µg) (C) Quantification of (B) showing the number of PKR-DRIF per field of view at the two concentrations. (D) IFA on A549 cells expressing GFP-PKR either mock treated or treated with Poly(I:C). GFP-PKR (green), p-eIF2α (magenta). The yellow arrows indicate cells that lack PKR-DRIF. The white arrows indicate cells containing PKR-DRIF. (E) Quantification of p-eIF2α in cells with or without PKR-DRIF. (F) Western blot confirming expression of GFP or GFP-NS4a into parental A549 cells. (G) IFA for PKR and dsRNA in A549 cells expressing either GFP or GFP-NS4a. White arrows indicate cells contain PKR-DRIF. Yellow arrows indicate cells with NS4a condensed on dsRNA and lacking PKR-DRIF. (H) Quantification of the percentage of cells per field of view (dot) expressing GFP or GFP-NS4a containing PKR-DRIF as represented in (G).

Because nsp15 and NS4a limit dsRNA loads and/or binding potential, we tested if increasing dsRNA loads promotes PKR-DRIF assembly by transfecting cells with either a low dose or a high dose of poly(I:C), which is a synthetic viral dsRNA mimic. These results showed that PKR-DRIF assembly significantly increased with the higher dose of poly(I:C) (Figure 5B,C and S8A), indicating that dsRNA loads positively correlate to PKR-DRIF assembly. To determine if poly(I:C)-induced PKR-DRIF assembly correlates to the activation of the PKR pathway, we generated an A549-PKR-KO cell line that stably expresses GFP-PKR (Figure S8B). We then lipofected cells with poly(I:C) and performed IFA for p-eIF2α. Importantly, cells containing PKR-DRIF had significantly higher p-eIF2α levels in comparison to cells lacking PKR-DRIF (Figure 5D,E). Notably, phosphorylation of eIF2α was dependent on GFP-PKR expression, as parental PKR-KO cells and PKR-KO cells expressing GFP-only control did not trigger the robust phosphorylation of eIF2α as observed in PKR-KO cells expressing GFP-PKR (Fig. S8C). These data indicate that the assembly of PKR-DRIF correlate to the activation of the PKR pathway.

We next asked if NS4a antagonizes PKR activation by competing with PKR for dsRNA-binding via condensation. To test this, we generated A549 cell lines that stably express either GFP or GFP-tagged NS4a (Figure S9A), which we verified by western blot (Figure 5F). We observed robust condensation of GFP-NS4a, but not the GFP-control, on poly(I:C) (Figure S9B). The condensation of GFP-NS4a on poly(I:C) resulted in a reduction in PKR condensation (Figure 5G,H). Importantly, NS4a displayed a higher partitioning coefficient on condensed dsRNA than PKR (Figure S9C,D), indicating that NS4a has a higher binding affinity for dsRNA and/or higher potential to condense on dsRNA relative to PKR.

### NS4a condensation antagonizes PKR activation in response to MERS-nsp15^mut^/ΔNS4a

In MERS-nsp15^mut^/ΔNS4a-infected cells, we observed GFP-NS4a but not GFP-control condense into puncta that co-localized with dsRNA within the ROR (Figure 6A,B). To test if NS4a condensation on viral dsRNA reduced the ability of PKR to activate, we performed IFA for p-PKR and dsRNA in cells following infection with MERS-nsp15^mut^/ΔNS4a. In comparison to the GFP-control cells, in which the majority of cells stained for p-PKR and contained PKR-DRIF, MERS-nsp15^mut^/ΔNS4a-infected cells expressing GFP-NS4a lacked p-PKR staining (Figure 6C,D). Moreover, the cells that lacked PKR-DRIF contained NS4a-dsRNA condensates (Figure 6C,E). Notably, the number of NS4a puncta were similar to the number of p-PKR puncta in GFP-control cells (Figure 6C,E). Importantly, expression of GFP-NS4a abolished the phosphorylation of eIF2α (Figure 6F,G), indicating that its ability to condense on dsRNA and prevent PKR condensation abolished the activation of the PKR pathway in response to MERS-nsp15^mut^/ΔNS4a.

**Figure 6.**
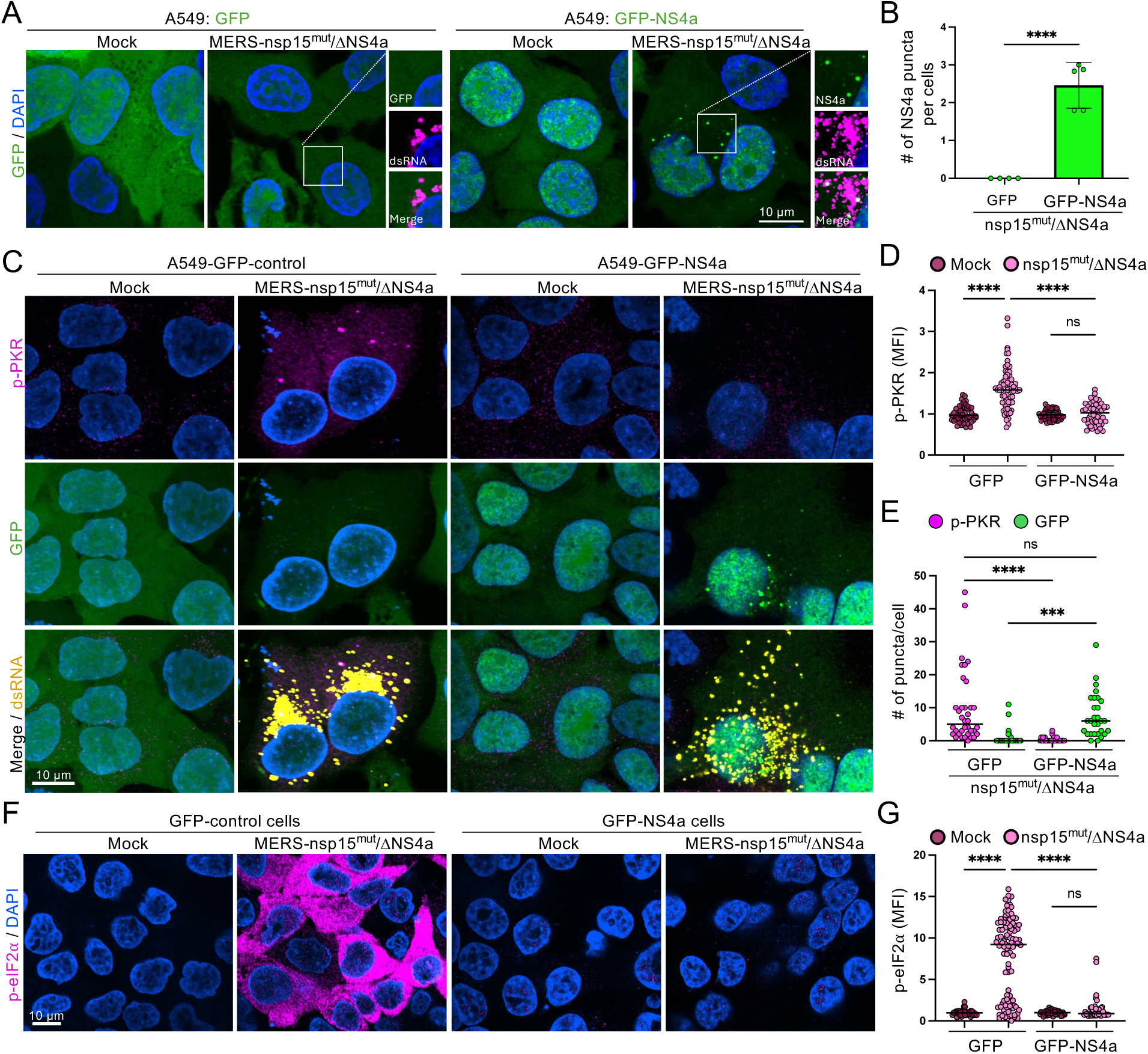
NS4a antagonizes PKR condensation and activation during MERS-CoV infection. (A) IFA for dsRNA (magenta) in A549-DPP4 cells expressing GFP (GFP-control) or GFP-NS4a 24 hrs. p.i. MERS-nsp15^mut^/ΔNS4a or mock-infected. (B) Quantification of (A) averaging the number of NS4a puncta per MERS-nsp15^mut^/ΔNS4a infected cell in both cells lines (each dot represents one field of view). (C) IFA for dsRNA (yellow) and p-PKR (magenta) in A549-DPP4 cells expressing GFP (GFP-control) or GFP-NS4a 24 hrs. p.i. MERS-nsp15^mut^/ΔNS4a or mock-infected. (D) Quantification of total cytosolic p-PKR mean fluorescence intensity (MFI) per cell (dot) as represented in (C). (E) Quantification of the number of GFP or p-PKR puncta in cells as represented in (C). (F) IFA for p-eIF2α in A549-DPP4 cells expressing GFP-control or GFP-NS4a following mock or MERS-nsp15^mut^/ΔNS4a infection. (G) Quantification of the mean fluorescence intensity (MFI) of p-eIF2α per cell (dot) as represented in (F).

These data demonstrate three important principles. First, NS4a condensation on viral dsRNA can prevent PKR condensation and activation. Second, NS4a forms puncta at only a select number of dsRNA foci in MERS-nsp15^mut^/ΔNS4a-infected cells, and this number is comparable to the number of PKR-DRIF observed in GFP-control cells. These observations further support that the viral dsRNA is exposed to the cytosol at select DMV-RTCs, which is where PKR can recognize and bind to the viral substrate. Third, the ability of NS4a to inhibit PKR condensation at these sites correlates with a reduction in PKR activation, further arguing that PKR condensation is a critical aspect of the PKR activation. Overall, these observations support that NS4a binds DMV-associated dsRNA during MERS-CoV infection, and that this limits PKR activation by occluding PKR from dsRNA exposed at select DMV-RTCs.

### PKR-DRIF correlate to PKR activation in response to Zika virus

To determine if PKR-DRIF formation is a more general occurrence in response to virus families other than coronaviruses, we infected A549 cells with Zika virus, an unrelated flavivirus, and performed IFA for PKR. In response to ZIKV infection, PKR assembled into puncta that typically co-localized with dsRNA (Figure 7A and Figure S11A). Greater than 70% of ZIKV-infected cells assembled these PKR puncta (Figure 7B), with each cell assembling an average of 10 PKR puncta (Figure 7C).

**Figure 7.**
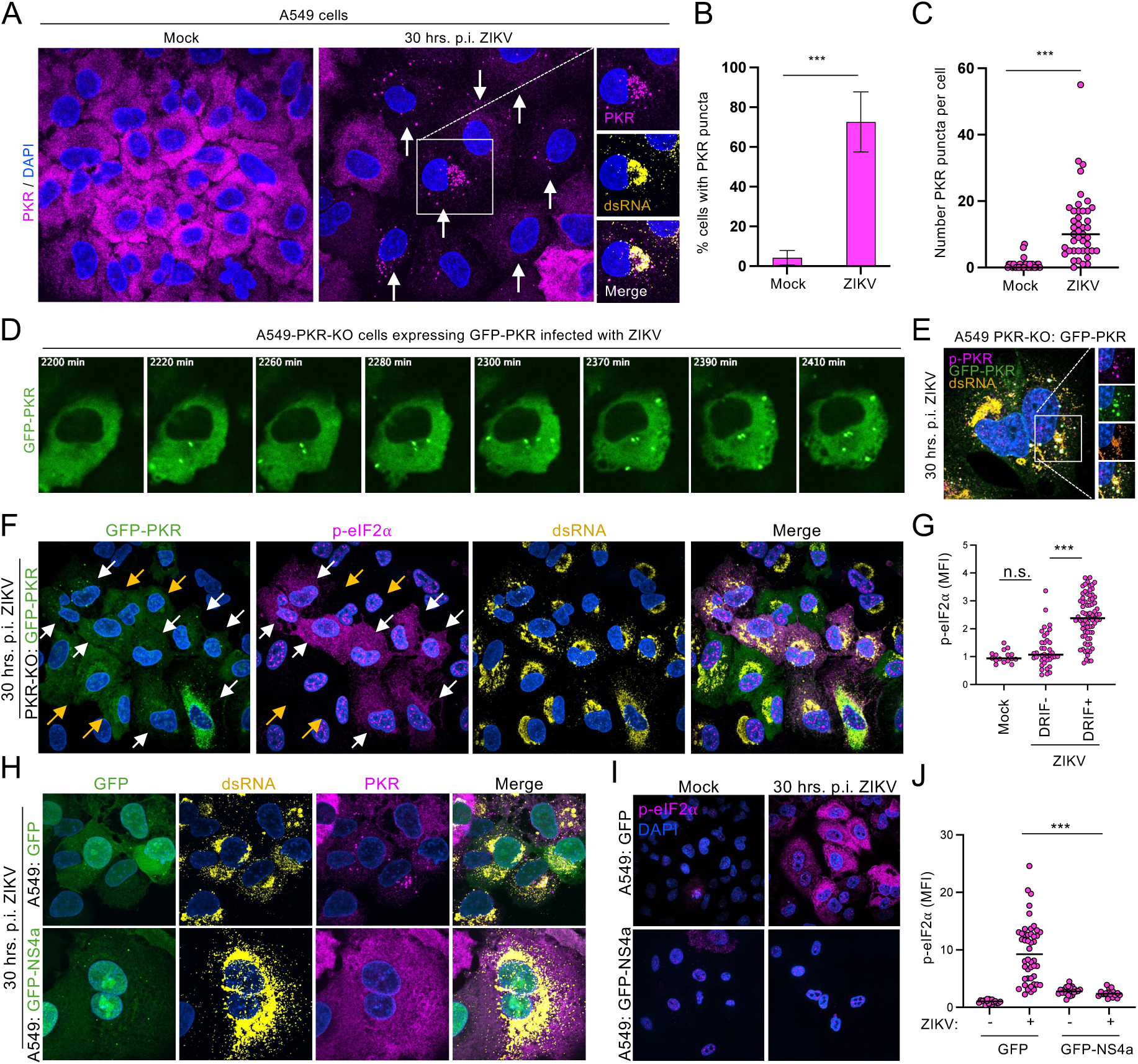
PKR-DRIF correlate to PKR activation in response to Zika virus and are inhibited by NS4a. (A) IFA for PKR and dsRNA in A549 cells 30 hrs. p.i. ZIKV infection (MOI=1) or mock treatment. The large panels show PKR staining. Arrows indicate cells containing PKR condensed on dsRNA. (B) Quantification of the percent of cells containing at least one PKR puncta as represented in (A). (C) Number of PKR puncta per cell (dot) as represented in (A). (D) Live cell imaging of A549 PKR-KO cells expressing GFP-PKR following ZIKV infection. (E) IFA for p-PKR (magenta) and dsRNA (yellow) in A549-PKR-KO cells expressing GFP-PKR. (F) IFA for p-eIF2α and dsRNA 30 hrs. p.i. in A549 PKR-KO cells expressing GFP-PKR. White arrows indicate cells with PKR-DRIF (DRIF+). Yellow arrows indicate cells lacking PKR-DRIF (DRIF-). (G) Quantification of the mean fluorescence intensity (MFI) of p-eIF2α per cell (dot) as represented in (F). (H) IFA for dsRNA and PKR in A549 cells expressing either GFP or GFP-NS4a 30 hrs. p.i. ZIKV or mock treatment. (I) IFA for p-eIF2α. (J) Quantification of the mean fluorescence intensity (MFI) of p-eIF2α per cell (dot) as represented in (I).

We further confirmed the assembly of PKR-DRIF by live-cell imaging of A549 PKR-KO cells stably expressing GFP-PKR. These analyses showed that PKR-DRIF assembly occurs between 12 and 24 hrs. p.i. ZIKV (Figure 7D and Supplemental Video S5 and S6). Notably, PKR-DRIF containing p-PKR assembled in perinuclear region of cells where ZIKV replication primarily occurs, and then re-localize to the cytosol (Figure 7E). These data suggest that the PKR-DRIF are sites of PKR activation in response to ZIKV infection.

We next tested if the formation of PKR-DRIF correlated to PKR-mediated phosphorylation of eIF2α. To do this, we performed IFA for p-eIF2α in PKR-KO cells expressing GFP-PKR. Notably, the expression of GFP-PKR in PKR-KO cells rescued the phosphorylation of eIF2α in response to ZIKV infection (Figure S11B-D), indicating that GFP-PKR is functional and that the phosphorylation of eIF2α in response to ZIKV is primarily mediated by PKR. Importantly, we observed that cells containing PKR-DRIF (DRIF+) had significantly higher levels of p-eIF2α in comparison to cells lacking PKR-DRIF (DRIF-) (Fig. 7F,G). These data argue that PKR-DRIF assembly coincides with the activation of PKR.

Further supporting that PKR-DRIF assembly correlates to PKR activation, we observed that MERS-CoV NS4a inhibited PKR-DRIF assembly in response to ZIKV infection (Fig. 7H). The inhibition of PKR-DRIF coincided with the formation of GFP-NS4a condensates within the ZIKV ROR (Figure 7H), suggesting that NS4a condensing on dsRNA prevents PKR binding to ZIKV dsRNA. The inhibition of PKR-DRIF by NS4a coincided with a significant reduction in p-eIF2α (Figure 7I,J), indicating that GFP-NS4a can inhibit PKR activation in response to ZIKV. Combined, these data further support that PKR-DRIF assemble is a vital process in the activation of PKR in response to ZIKV.

## DISCUSSION

We propose a model for PKR activation based on our microscopy analyses of PKR activation in response to MERS-nsp15^mut^/ΔNS4a and ZIKV (Figure 8). In this model, PKR condenses on dsRNA exposed at select viral DMVs and undergoes autophosphorylation. Activated p-PKR then forms condensates termed PKR-<u>d</u>s<u>R</u>NA induced foci (DRIF). PKR-DRIF disassociate from dsRNA and subsequently disassemble, which releases p-PKR molecules into the cytosol where they phosphorylate eIF2α to initiate the ISR and arrest protein synthesis. An important implication of this model is that the dissociation of PKR-DRIF from dsRNA allows for additional cycles of inactive PKR monomers to bind the same dsRNA substrates and undergo activation, which could be critical for allowing robust activation of PKR from limited dsRNA substrates exposed from only a small fraction of viral DMVs.

**Figure 8.**
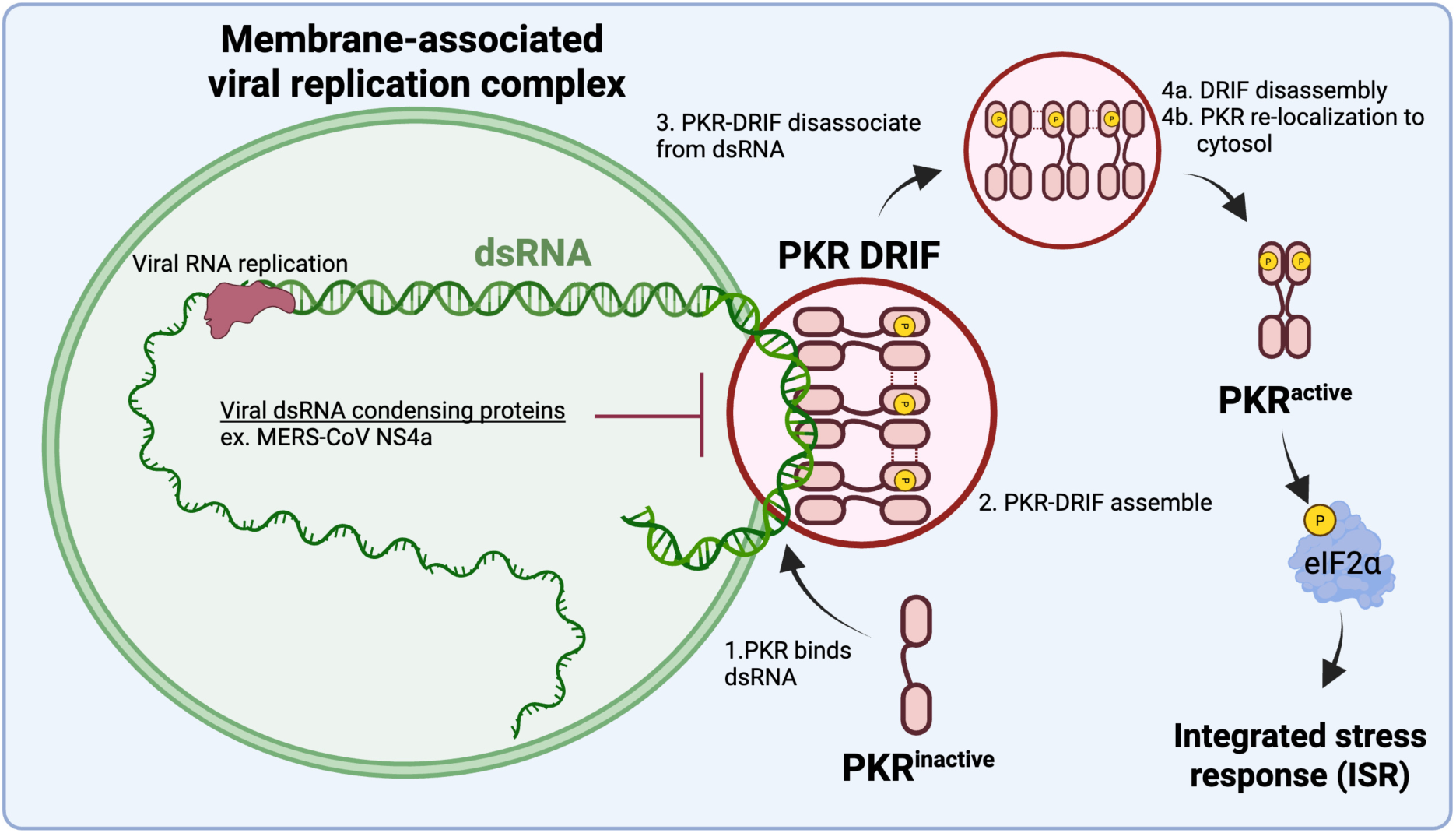
Model of PKR-DRIF functions during PKR activation in response viruses. Monomeric inactive PKR binds to dsRNA exposed from membrane-associated viral RTCs. PKR condenses on dsRNA, leading to PKR activation and the formation of PKR-DRIF. PKR-DRIF disassociate from viral dsRNA and disassemble, releasing p-PKR molecules into the cytosol where they phosphorylate their substrates. Additional monomeric inactive PKR binds to the exposed dsRNA, creating an activation cycle.

The following key observations support this model. The notion that PKR condenses and activates on dsRNA associated with viral DMVs is supported by the observations that: i) PKR-DRIF are the initial sites of PKR activation (Figure 2E); ii) PKR-DRIF are the primary sites of interaction between of p-PKR and dsRNA (Figure 2F); iii) PKR-DRIF are composed of p-PKR oligomers (3-30 molecules) located on the cytosolic surface of viral DMVs (Fig, 2G-I). The notion that PKR-DRIF disassociate from dsRNA/DMVs and disassemble is based on the observations that: i) some PKR-DRIF did not co-localize with viral dsRNA/DMV, yet they were invariably localized near viral dsRNA/DMVs (Figures 2G and 3A); ii) PKR-DRIF were rarely observed outside the ROR (Figure 3B); iii) PKR-DRIF underwent fission and dissolution (Figure 3E). The data supporting that p-PKR molecules are predominantly localized in the cytosol and disassociated from dsRNA is that p-PKR diffusely localized in the cytosol did not co-localize with dsRNA by IFA and lacked detectable interaction with dsRNA by PLA (Figures 1A and 2A,F). The notion that p-PKR re-localizes from the ROR to the cytosol is based on the observations that: i) p-PKR levels in the cytosol lagged behind p-PKR levels in the ROR (Figures 4A,B); ii) PKR-mediated phosphorylation of eIF2α primarily occurred in the cytosol and not in the ROR (Figures 4C,D).

The observation that most viral DMVs were not associated with PKR-DRIF (Figures 2A-I) suggests that only a small fraction of viral DMVs trigger PKR activation. We propose three possibilities that explain why select DMVs trigger PKR-DRIF assembly. First, these DMVs could represent a transient morphological state that is prone to exposing dsRNA to the cytosol, such as DMV fusion, which has been shown to take place during coronavirus infection ^38,39^. Second, these DMVs may become damaged during the course of viral replication, leading to dsRNA exposure to the cytosol. Third, a small fraction of DMVs could export the negative strand, thus leading to dsRNA formation immediately upon its export to the cytosol where positive strands are abundant. Regardless of the cause, the high frequency of cells that activate PKR in response to MERS-nsp15^mut^/ΔNS4a demonstrates that a critical fraction of DMVs do not completely protect viral RTC from PRR sensing during MERS-CoV infection, thus necessitating MERS-CoV to encode nsp15 and NS4a to prevent detection of dsRNA by PKR. Ongoing studies are investigating the morphological alterations of the select DMVs that trigger PKR activation.

Previous analyses of PKR localization during the antiviral response showed that PKR localizes to stress granules ^30,31^. However, subsequent studies did not corroborate this finding ^27,28^. Instead, PKR was shown to assemble into higher-order structures composed of dsRNA-binding proteins and dsRNA that were distinct from stress granules. These structures were termed DRIF and were proposed to initiate PKR activation in response to synthetic dsRNA^27^. Our data further supports that PKR assembles into DRIF in response to viral infection, and that this coincides with the activation of PKR. Moreover, we show that PKR-DRIF represent the primary sites of interactions between PKR and dsRNA during viral infection. Lastly, our data support a novel function of PKR-DRIF, whereby PKR-DRIF promote disassociate of p-PKR from viral dsRNA, thus becoming a dsRNA-less PKR-DRIF that then disassembles (Figures 2A,B). Therefore, we propose two functions of PKR-DRIF. First, condensation of PKR into a DRIF promotes dimerization and autophosphorylation of PKR. Second, PKR-DRIF then promote the disassociation via a yet-to-be-determined mechanism, thus allowing for activated p-PKR to re-localize to the cytosol where its substrates are located.

An important implication of our data is that it supports that activated PKR disassociates from dsRNA. This is based on the following key observations: i) dsRNA was not diffusely localized to the cytosol where p-PKR molecules localize as assessed by IFA and immunogold TEM (Figures 1A and 2A). Instead, dsRNA primarily localized to DMVs within the ROR, which is where it interacted with PKR-DRIF (Fig. 2); ii) p-PKR molecules diffusely localized in the cytosol did not display interaction with dsRNA by PLA or co-localize with dsRNA by IFA (Fig. 2). However, we do not rule out the possibility that viral dsRNA is fragmented upon PKR-DRIF assembly, resulting in p-PKR molecules in the cytosol that are bound to small fragments of dsRNA unable to be detected by the K1 dsRNA antibody used in our PLA, TEM, and IFA analyses. Notably, the K1 dsRNA antibody is able to bind dsRNA as short as 40 base pairs, which is just slightly longer than the minimal 30 base pairs required for promoting PKR dimerization ^40–42^, thus making this scenario unlikely but nevertheless possible. The ability of PKR to remain in an activated state after dsRNA disassociation is likely dependent on the catalytic domain remaining phosphorylated. Future studies will address the mechanisms that underly p-PKR disassociation from dsRNA and the kinetics of p-PKR dephosphorylation upon dsRNA disassociation.

How dsRNA-less PKR-DRIF maintain their structure once disassociated from dsRNA is not understood. Based on our immunogold TEM for p-PKR (Figure 2I), we propose that p-PKR dimers could oligomerize and/or form condensates via weak multivalent interactions. This process could destabilize interactions with dsRNA to promote disassociation of PKR-DRIF from dsRNA, though additional structural studies on PKR will be required to test this hypothesis. However, we do not rule out that additional cellular factors could scaffold p-PKR into a condensate or that cellular (helicases) or viral factors [RdRP; helicase (nsp8)] could be responsible for causing PKR-DRIF disassociation from dsRNA. Regardless, our data suggest that PKR-DRIF readily disassemble upon disassociation from dsRNA (Figure 3D). Future work will aim to understand the assembly, maintenance, and disassembly of PKR-DRIF.

The elevated levels of dsRNA observed in cells infected with MERS-nsp15^mut^ in comparison to parental MERS-WT suggest that MERS-CoV nsp15 limits dsRNA accumulation (Figure 5A), which supports previous data ^7^. This is also consistent with the function of Endo U encoded by SARS-CoV-2 and MHV ^8,10,43^. However, we observed an additional elevation in IFA signal for dsRNA in response to the MERS-ΔNS4a and MERS-nsp15^mut^/ΔNS4a (Figure 5A), which we reason is due to enhanced dsRNA binding by the K1 antibody because structural and *in vitro* studies have indicated that NS4a is a dsRNA-binding protein ^7,14^. These data indicate that while EndoU and NS4a each alone reduce PKR activation to some extent, the combination of both EndoU and NS4a robustly shut down the PKR pathway (Figure 1). Consistent with this, we observed that NS4a can robustly condense on both synthetic dsRNA and viral dsRNA (Figures 5B,C and 6G). Importantly, our data support that NS4a condensation on dsRNA inhibits PKR condensation and activation during poly(I:C) treatment (Figures 5G,H), MERS-nsp15^mut^/ΔNS4a infection (Fig. 6C-G), or Zika virus infection (Figures 7H-J). Combined, these data support that NS4a condenses on dsRNA exposed from DMVs to limit the ability of PKR to condense and activate. Future work will investigate the properties of NS4a that allow it to have a competitive advantage in dsRNA condensation relative to PKR.

In conclusion, we showed that PKR condensation on viral dsRNA exposed to the outer surface of DMVs initiates the PKR pathway, and that MERS-CoV NS4a directly antagonizes this process. This establishes that a host-virus arms race centers on the dsRNA condensation potential of PRRs and viral proteins. It will be imperative to further characterize the process of PKR condensation, examine the subcellular localization of PKR activation during more diverse viral infections (DNA viruses, negative strand RNA viruses), and examine the effect of NS4a condensation on dsRNA on additional innate immune pathways.

## ACKNOWLEDGMENTS

Transmission electron microscopy was overseen by Dr. Naomi Kamasawa at the electron microscopy core at the Max Planck Florida Institute for Neuroscience. This work was funded by institutional funds from the Herbert Wertheim University of Florida Scripps Institute for Biomedical Innovation and Technology (JMB), the National Institutes of Health (NIH) awards R35GM151249 (JMB), R01AI140442 (SRW) and R01AI169537 (SRW). NB was supported in part by NIH K12GM081259.

## AUTHOR CONTRIBUTIONS

JMB, SRW, EKB, NB conceived the project. JMB, EKB, NB, HW, MJW performed experiments. JMB, SRW, EKB, and NB analyzed data. EKB and JMB wrote manuscript. JMB, NB, SWR, and EKB edited and revised manuscript.

## COMPETING INTERESTS

The authors declare no competing interests

## MATERIALS & METHODS

### Cell lines

A549 cells or A549 cells expressing the MERS-CoV receptor, DPP4 ^7^, were used as the primary cell line, unless indicated. Additional cell lines utilized include are described in Supplemental spreadsheet. All A549 cells were cultured in RPMI with 1x Penicillin/Streptomycin and 10% heat-inactivated FBS. Vero CCL-81 cells were used for viral quantification and cultured in DMEM supplemented with 1x Penicillin/Streptomycin and 10% heat-inactivated FBS. All cells were cultured at 37°C with 5% CO_2_.

For the cell lines generated in preparation of this manuscript, with indicated source being “this work” in Supplemental spreadsheet, lentiviruses were made as described in ^44^. In brief, human embryonic kidney–293T cells (T-25 flask; 80% confluent) were co-transfected with 2.4 μg of lentiviral transfer plasmid (pLenti-EF1-Blast^R^), 0.8 μg of pVSV-G, 0.8 μg of pRSV-Rev, and 1.4 μg of pMDLg-pRRE using 15 μl of Lipofectamine 2000. Medium was collected at 48 hours post transfection and filter-sterilized with a 0.45μm filter. A549 cells were transduced as described in ^45^. Briefly, medium was removed from A549 cells (T-25 flask; 80% confluent), before 1ml lentivirus was added to the flasks and left to incubate at 37°C with 5% CO_2_ for 1h with occasional rocking. After 1 hour of incubation, fresh medium was added. The medium was changed 24 hours post transduction, and placed on selection medium 72 hours post transduction. The selection medium was changed every 3 days. After 1 week, the selection medium was replaced with a normal growth medium. Protein expression was confirmed by immunoblot analysis.

### Plasmid generation

To generate GFP-tagged PKR plasmid, 5’-GFP-tagged PKR coding sequence was synthesized as a gBlock by IDT (Supplemental spreadsheet) and assembled into pLenti-EF1-Blast vector. In more detail, GFP-PKR was inserted at the XhoI/XbaI restriction sites using Gibson assembly. For the GFP-NS4a plasmids, the PKR CDS was removed from the pLenti-EF1-Blast-GFP-PKR plasmid (previously described) using the restriction sites of XbaI flanking the PKR CDS. The NS4a sequence containing overhangs complementary to the vector was synthesized as a gBlock (Supplemental spreadsheet), and inserted into the pLenti-EF1-Blast-GFP vector using Gibson assembly.

### Viral infections

For MERS-CoV infections, the designated type of A549-DPP4 cells were mock infected or infected with either MERS-WT, MERS-nsp15^mut^, or MERS-ΔNS4a, MERS-nsp15^mut^/NS4a ^7^. All infections were performed at an MOI =1 PFU/cell, unless otherwise stated, with a 1hr adsorption period at 37°C. After adsorption, cells were washed with 1x PBS, and RPMI with 2% heat-inactivated FBS was added. Cells were incubated at 37 °C until the indicated time post-infection. If translation activity was to be examined, cells were treated with 10μg/ml of puromycin for 10 minutes prior to collection. All infections and virus manipulations were carried out in a biosafety level 3 laboratory (BSL-3) utilizing appropriate personal protective equipment as approved by the University of Pennsylvania Institutional Biosafety Committee (IBC). For Zika virus (ZIKV) infections, indicated A549 cell lines were either mock infected or infected with ZIKV (MOI 5) by incubating cells (12-well format) with 0.5 ml of FBS-free medium containing virus for 2 hours with periodic rocking. Normal growth medium was added after 2 hours, and cells were fixed at times stated in the figures. Stock of Zika virus (PB 81) was generated in Vero cells, and titer was determined by plaque assay on Vero cells. All viral infections were performed in BSL2+ laboratory at the Herbert Wertheim UF Scripps Institute for Biomedical Innovation and Technology.

### Plaque assay

Supernatants from infected cells were collected at the indicated time point post-infection. Infectious virus was quantified via plaque assay on Vero CCL-81 cells as described previously^8^.

### Poly(I:C) treatment

For each well on a 12-well plate, 500ng Poly(I:C) was combined with 1.5 μl of Lipofectamine 2000 in 100 μl serum-free DMEM. The mixture was left to incubate at room temperature for 15 minutes, before being added to A549 cells at 80% confluency for time indicated in the figures. Thereafter, coverslips would be prepared accordingly to the IFA methods previously described, or lysates would be collected before following the method for western blotting (explained below).

### Western blot analysis

Cell lysates were harvested at the indicated time points post-infection using a lysis buffer as previously described ^8^. Lysates were mixed with 4x Laemmli sample buffer 1x DTT (Dithiothreitol), and heated to 95 °C for 10 min. Samples were resolved on gradient SDS/PAGE gels (Bio-Rad) and transferred to polyvinylidene difluoride (PVDF) membrane. Membranes were blocked with either 5% bovine serum albumin (BSA) or 5% non-fat milk in Tris-buffered saline with 0.1% Tween (TBST) and then probed with the indicated primary antibody overnight at 4 °C (Supplemental spreadsheet). Blots were washed and subsequently probed with the designated secondary antibody for 1 hour at room temperature. Blots were visualized using SuperSignal West Femto Chemiluminescent Substrate.

### Live cell imaging

A549-DPP4 +GFP-PKR were either mock-infected or infected, as described above, with MERS-WT or MERS-nsp15^mut^/NS4a at an MOI of 5 in a glass-bottom 96-well plate. After infection cells were incubated until ∼12 hours post-infection and then placed in the live cell imaging chamber of the Molecular Devices Image Xpress instrument. The chamber was kept at 37 °C and 5% CO2. Images of the FITC channel were taken ∼10 to 20 minutes over 48 hours. Infections and image acquisition were performed in BSL-3 in accordance with University of Pennsylvania Institutional Biosafety Committee (IBC) approved protocols.

For ZIKV live cell imaging, A549-PKR-KO cells expressing GFP-PKR were seeded on glass bottom dishes. Cells were infected at an MOI=1 and imaged via confocal microscopy. Cells were maintained in an environmental chamber at 37 degrees Celsius and 5% CO_2_ throughout imaging.

### Immunofluorescence Assay (IFA) and smRNA-FISH

The indicated A549 cells type was seeded onto glass coverslips (Thomas Scientific: 1203J81) and infected as described above. At the indicated time point, samples were fixed with 4% paraformaldehyde (PFA) for 30 min prior followed by permeabilization in 70% ethanol. For IFA, the samples were incubating with the indicated primary antibody diluted in PBS overnight at 4C (Supplemental spreadsheet). All secondary antibodies were Alexa Fluor antibodies purchased from Abcam and added at 1:1,000 in PBS. Cells were washed 3x in PBS, then secondary antibodies were diluted in PBS and added for 2 h at room temperature. Coverslips were then washed 3x in PBS before being mounted on coverslides (Fisher: 12-544-11) with Vectashield (Vector Laboratories: 101098-044) or ProLong™ Gold Antifade Mountant (Invitrogen: P36931). For dual staining of IFA and smFISH, after washing off the secondary antibodies with PBS, the cells were fixed in 4% PFA (Fisher Scientific Co LLC: 50980495) for 10 min. Cells were washed 3x in PBS, before being placed in buffer A (filter-sterilized 2x SSC with 10% formamide) for 5 min. For each sample, 50uL of smFISH probes diluted 1:100 in hybridization buffer ((Fisher Scientific Co LLC: 09-719D), 10% dextran sulfate (Fisher Scientific Co LLC: S4030), 10% formamide (Fisher Scientific Co LLC: BP227500), 1x nuclease-free SSC (Life Technologies Corporation: 15557044), diluted in nuclease-free water (Fisher Scientific Co LLC: 10977023)) was prepared. The mixture was added into a petri dish (Fisher: FB0875711A), and the coverslips were then flipped onto the smFISH probes. Petri dishes were sealed with parafilm and incubated overnight at 37C. The next day, the coverslips were washed 2x in Buffer A, then once in 2x SSC. The coverslips were then mounted on coverslides in the same manner as described for IFA. Images were taken on a Nikon Eclipse Ti2 with a CFI60 Plan Apochromat Lambda D 100x Oil Immersion Objective Lens, N.A. 1.45, W.D. 0.13mm, F.O.V. 25mm, DIC, Spring Loaded. The filter set included: C-FL DAPI Filter Set, High-Signal-Noise, Semrock Brightline, Excitation: 356/30nm (341-371nm). Super-resolution microscopy was imaged using GATACA Systems Live-SR. Image processing and analysis was done using FIJI 2.16.0.

### smFISH probe labeling

MERS-ORF1a probes were generated using Stellaris probe design tool and stated in Supplemental spreadsheet. The smFISH probes were labeled with Atto-550 or Atto-633 using 5-Propargylamino-ddUTP-ATTO-550 (Jena Bioscience: NU-1619-550) or 5-Propargylamino-ddUTP-ATTO-633 (Jena Bioscience: NU-1619-633. Oligos were labeled by mixing 16 μM oligos, 2 μl of ddNTPs, 1x TDT buffer, and 1.5 μl of terminal deoxynucleotidyl transferase (Thermo Fisher Scientific: EP0161) together in a 50 μl total reaction volume. Oligos were then incubated overnight at 37C, and precipitated with 400 ml ethanol and 50 ml 3M Sodium acetate (pH 3.2) at -20C overnight. The precipitated oligos were spun down at 20,000xg for 30 min, the supernatant was aspirated, and the pellet was washed twice with 75% ethanol. After the second wash, the pellets were left to dry and then resuspended into 80 ml RNase free water.

### Proximity Ligation Assay (PLA)

Proximity ligation assay (Millipore Sigma: DUO92101-1KT) was performed according to manufacturer’s protocol (Millipore Sigma). In brief, coverslips were incubating with blocking buffer for 1 hour at 37°C, before primary antibodies were added in the same manner as during IFA but diluted in provided PLA antibody dilutant. The coverslips were washed 2x with Wash Buffer A, before adding the plus and minus probe provided in the PLA kit. In instances where co-IF was conducted, the coverslips were incubated with the probe and AlexaFluor secondary antibodies simultaneously. The samples were placed in the 37°C incubator for 60 min, before being washed 2x with Wash Buffer A. After, the coverslips were incubated with the ligation reaction for 30 min at 37°C before washing 2x with Wash Buffer A. Lastly, the coverslips were left to incubate with the polymerase reaction at 37°C for 100min, before being washed 2x with Wash Buffer B and left in 0.01x Wash Buffer B. The samples were mounted on slides with mounting media containing DAPI before being imaged.

### Transmission Electron Microscopy (TEM) sample preparation

A549-DPP4 cells were seeded onto plastic coverslips (Electron Microscopy Sciences: 72280) and infected as described above. At the indicated time point post infection, half of the media was removed and replaced with a fixing buffer composed of 4% PFA, 0.1% glutaraldehyde, and 0.1M Sorenson’s buffer. After 5 minutes the entire volume was removed and replaced with fresh fixing buffer for 30 minutes. Samples were washed three times and stored in 0.1M Sorenson’s buffer overnight. The next day, the samples were placed in 15% sucrose solution diluted in 0.1M Sorenson’s buffer for 1 hour at 4C, before being placed in 30% sucrose solution diluted in Sorenson’s buffer for another 2 hours at 4C. Thereafter, the coverslips were plunged into liquid nitrogen for 1 min before thawing at room temperature. This was repeated one more time, before placing the samples into 0.1M Sorenson’s buffer. Finally, the coverslips were placed in primary antibody solution containing either anti-p-PKR (1:1,000) or anti-dsRNA (1:1,000) antibodies diluted in 0.1M Sorenson’s buffer and placed at 4C for two nights. The following steps and imaging were conducted by Max Planck Florida Institute for Neuroscience (MPFI) Imaging Center.

### Quantification and statistical analysis

Image processing was conducted in Fiji (ImageJ2) and data processing was conducted in Microsoft Excel. Graphs and p-values were derived by student’s t-test (two variables tested) or two-way ANOVA (more than two variables tested) in Prism GraphPad. *p < 0.05, **p > 0.01, ***p > 0.001 unless otherwise noted in the figure legend. Error bars represent standard deviation, calculated by Prism GraphPad.

## Notes

### Competing Interest Statement

The authors have declared no competing interest.

## REFERENCES

1. Zhang, A.-R., Shi, W.-Q., Liu, K., Li, X.-L., Liu, M.-J., Zhang, W.-H., Zhao, G.-P., Chen, J.-J., Zhang, X.-A., Miao, D., et al. (2021). Epidemiology and evolution of Middle East respiratory syndrome coronavirus, 2012-2020. Infect. Dis. Poverty 10, 66. 10.1186/s40249-021-00853-0.

2. Chandran, D., Chakraborty, S., Chandran, D., Subedi, D., Jisha, A.I., Chopra, H., Rabaan, A.A., Al-Tawfiq, J.A., Islam, M.R., and Dhama, K. (2024). Middle east respiratory syndrome coronavirus could be a priority pathogen to cause public health emergency: noticeable features and counteractive measures. Environ. Health Insights 18, 11786302241271544. 10.1177/11786302241271545.

3. Otter, C.J., Renner, D.M., Fausto, A., Tan, L.H., Cohen, N.A., and Weiss, S.R. (2024). Interferon signaling in the nasal epithelium distinguishes among lethal and common cold coronaviruses and mediates viral clearance. Proc Natl Acad Sci USA 121, e2402540121. 10.1073/pnas.2402540121.

4. Tanneti, N.S., Stillwell, H.A., and Weiss, S.R. (2025). Human coronaviruses: activation and antagonism of innate immune responses. Microbiol. Mol. Biol. Rev. 89, e0001623. 10.1128/mmbr.00016-23.

5. Li, Y.-H., Hu, C.-Y., Wu, N.-P., Yao, H.-P., and Li, L.-J. (2019). Molecular Characteristics, Functions, and Related Pathogenicity of MERS-CoV Proteins. Engineering (Beijing) 5, 940–947. 10.1016/j.eng.2018.11.035.

6. Mihelc, E.M., Baker, S.C., and Lanman, J.K. (2021). Coronavirus infection induces progressive restructuring of the endoplasmic reticulum involving the formation and degradation of double membrane vesicles. Virology 556, 9–22. 10.1016/j.virol.2020.12.007.

7. Comar, C.E., Otter, C.J., Pfannenstiel, J., Doerger, E., Renner, D.M., Tan, L.H., Perlman, S., Cohen, N.A., Fehr, A.R., and Weiss, S.R. (2022). MERS-CoV endoribonuclease and accessory proteins jointly evade host innate immunity during infection of lung and nasal epithelial cells. Proc Natl Acad Sci USA 119, e2123208119. 10.1073/pnas.2123208119.

8. Otter, C.J., Bracci, N., Parenti, N.A., Ye, C., Asthana, A., Blomqvist, E.K., Tan, L.H., Pfannenstiel, J.J., Jackson, N., Fehr, A.R., et al. (2024). SARS-CoV-2 nsp15 endoribonuclease antagonizes dsRNA-induced antiviral signaling. Proc Natl Acad Sci USA 121, e2320194121. 10.1073/pnas.2320194121.

9. Ancar, R., Li, Y., Kindler, E., Cooper, D.A., Ransom, M., Thiel, V., Weiss, S.R., Hesselberth, J.R., and Barton, D.J. (2020). Physiologic RNA targets and refined sequence specificity of coronavirus EndoU. RNA 26, 1976–1999. 10.1261/rna.076604.120.

10. Hackbart, M., Deng, X., and Baker, S.C. (2020). Coronavirus endoribonuclease targets viral polyuridine sequences to evade activating host sensors. Proc Natl Acad Sci USA 117, 8094–8103. 10.1073/pnas.1921485117.

11. Frazier, M.N., Wilson, I.M., Krahn, J.M., Butay, K.J., Dillard, L.B., Borgnia, M.J., and Stanley, R.E. (2022). Flipped over U: structural basis for dsRNA cleavage by the SARS-CoV-2 endoribonuclease. Nucleic Acids Res. 50, 8290–8301. 10.1093/nar/gkac589.

12. Salukhe, I., Choi, R., Van Voorhis, W., Barrett, L., and Hyde, J. (2023). Regulation of coronavirus nsp15 cleavage specificity by RNA structure. PLoS ONE 18, e0290675. 10.1371/journal.pone.0290675.

13. Wright, Z.M., Butay, K.J., Krahn, J.M., Wilson, I.M., Gabel, S.A., DeRose, E.F., Hissein, I.S., Williams, J.G., Borgnia, M.J., Frazier, M.N., et al. (2025). Spontaneous base flipping helps drive Nsp15’s preferences in double stranded RNA substrates. Nat. Commun. 16, 391. 10.1038/s41467-024-55682-0.

14. Niemeyer, D., Zillinger, T., Muth, D., Zielecki, F., Horvath, G., Suliman, T., Barchet, W., Weber, F., Drosten, C., and Müller, M.A. (2013). Middle East respiratory syndrome coronavirus accessory protein 4a is a type I interferon antagonist. J. Virol. 87, 12489– 12495. 10.1128/JVI.01845-13.

15. Meurs, E., Chong, K., Galabru, J., Thomas, N.S., Kerr, I.M., Williams, B.R., and Hovanessian, A.G. (1990). Molecular cloning and characterization of the human double-stranded RNA-activated protein kinase induced by interferon. Cell 62, 379–390. 10.1016/0092-8674(90)90374-n.

16. Nanduri, S., Rahman, F., Williams, B.R., and Qin, J. (2000). A dynamically tuned double-stranded RNA binding mechanism for the activation of antiviral kinase PKR. EMBO J. 19, 5567–5574. 10.1093/emboj/19.20.5567.

17. Ung, T.L., Cao, C., Lu, J., Ozato, K., and Dever, T.E. (2001). Heterologous dimerization domains functionally substitute for the double-stranded RNA binding domains of the kinase PKR. EMBO J. 20, 3728–3737. 10.1093/emboj/20.14.3728.

18. Zhang, F., Romano, P.R., Nagamura-Inoue, T., Tian, B., Dever, T.E., Mathews, M.B., Ozato, K., and Hinnebusch, A.G. (2001). Binding of double-stranded RNA to protein kinase PKR is required for dimerization and promotes critical autophosphorylation events in the activation loop. J. Biol. Chem. 276, 24946–24958. 10.1074/jbc.M102108200.

19. Su, Q., Wang, S., Baltzis, D., Qu, L.-K., Wong, A.H.-T., and Koromilas, A.E. (2006). Tyrosine phosphorylation acts as a molecular switch to full-scale activation of the eIF2alpha RNA-dependent protein kinase. Proc Natl Acad Sci USA 103, 63–68. 10.1073/pnas.0508207103.

20. Koromilas, A.E., Roy, S., Barber, G.N., Katze, M.G., and Sonenberg, N. (1992). Malignant transformation by a mutant of the IFN-inducible dsRNA-dependent protein kinase. Science 257, 1685–1689. 10.1126/science.1382315.

21. Williams, B.R.G. (1995). The role of the dsRNA-activated kinase, PKR, in signal transduction. Seminars in virology 6, 191–202. 10.1006/smvy.1995.0024.

22. Donnelly, N., Gorman, A.M., Gupta, S., and Samali, A. (2013). The eIF2α kinases: their structures and functions. Cell. Mol. Life Sci. 70, 3493–3511. 10.1007/s00018-012-1252-6.

23. Gordiyenko, Y., Llácer, J.L., and Ramakrishnan, V. (2019). Structural basis for the inhibition of translation through eIF2α phosphorylation. Nat. Commun. 10, 2640. 10.1038/s41467-019-10606-1.

24. Costa-Mattioli, M., and Walter, P. (2020). The integrated stress response: From mechanism to disease. Science 368. 10.1126/science.aat5314.

25. Dalet, A., Gatti, E., and Pierre, P. (2015). Integration of PKR-dependent translation inhibition with innate immunity is required for a coordinated anti-viral response. FEBS Lett. 589, 1539–1545. 10.1016/j.febslet.2015.05.006.

26. Rabouw, H.H., Langereis, M.A., Knaap, R.C.M., Dalebout, T.J., Canton, J., Sola, I., Enjuanes, L., Bredenbeek, P.J., Kikkert, M., de Groot, R.J., et al. (2016). Middle East Respiratory Coronavirus Accessory Protein 4a Inhibits PKR-Mediated Antiviral Stress Responses. PLoS Pathog. 12, e1005982. 10.1371/journal.ppat.1005982.

27. Corbet, G.A., Burke, J.M., Bublitz, G.R., Tay, J.W., and Parker, R. (2022). dsRNA-induced condensation of antiviral proteins modulates PKR activity. Proc Natl Acad Sci USA 119, e2204235119. 10.1073/pnas.2204235119.

28. Cusic, R., and Burke, J.M. (2024). Condensation of RNase L promotes its rapid activation in response to viral infection in mammalian cells. Sci. Signal. 17, eadi9844. 10.1126/scisignal.adi9844.

29. Zappa, F., Muniozguren, N.L., Wilson, M.Z., Costello, M.S., Ponce-Rojas, J.C., and Acosta-Alvear, D. (2022). Signaling by the integrated stress response kinase PKR is fine-tuned by dynamic clustering. J. Cell Biol. 221. 10.1083/jcb.202111100.

30. Reineke, L.C., Kedersha, N., Langereis, M.A., van Kuppeveld, F.J.M., and Lloyd, R.E. (2015). Stress granules regulate double-stranded RNA-dependent protein kinase activation through a complex containing G3BP1 and Caprin1. MBio 6, e02486. 10.1128/mBio.02486-14.

31. Paget, M., Cadena, C., Ahmad, S., Wang, H.-T., Jordan, T.X., Kim, E., Koo, B., Lyons, S.M., Ivanov, P., tenOever, B., et al. (2023). Stress granules are shock absorbers that prevent excessive innate immune responses to dsRNA. Mol. Cell 83, 1180–1196.e8. 10.1016/j.molcel.2023.03.010.

32. Comar, C.E., Goldstein, S.A., Li, Y., Yount, B., Baric, R.S., and Weiss, S.R. (2019). Antagonism of dsRNA-Induced Innate Immune Pathways by NS4a and NS4b Accessory Proteins during MERS Coronavirus Infection. MBio 10. 10.1128/mBio.00319-19.

33. te Velthuis, A.J.W., van den Worm, S.H.E., and Snijder, E.J. (2012). The SARS-coronavirus nsp7+nsp8 complex is a unique multimeric RNA polymerase capable of both de novo initiation and primer extension. Nucleic Acids Res. 40, 1737–1747. 10.1093/nar/gkr893.

34. Snijder, E.J., Limpens, R.W.A.L., de Wilde, A.H., de Jong, A.W.M., Zevenhoven-Dobbe, J.C., Maier, H.J., Faas, F.F.G.A., Koster, A.J., and Bárcena, M. (2020). A unifying structural and functional model of the coronavirus replication organelle: Tracking down RNA synthesis. PLoS Biol. 18, e3000715. 10.1371/journal.pbio.3000715.

35. de Wilde, A.H., Raj, V.S., Oudshoorn, D., Bestebroer, T.M., van Nieuwkoop, S., Limpens, R.W.A.L., Posthuma, C.C., van der Meer, Y., Bárcena, M., Haagmans, B.L., et al. (2013). MERS-coronavirus replication induces severe in vitro cytopathology and is strongly inhibited by cyclosporin A or interferon-α treatment. J. Gen. Virol. 94, 1749– 1760. 10.1099/vir.0.052910-0.

36. Watkins, J.M., and Burke, J.M. (2024). A closer look at mammalian antiviral condensates. Biochem. Soc. Trans. 52, 1393–1404. 10.1042/BST20231296.

37. Deng, X., Hackbart, M., Mettelman, R.C., O’Brien, A., Mielech, A.M., Yi, G., Kao, C.C., and Baker, S.C. (2017). Coronavirus nonstructural protein 15 mediates evasion of dsRNA sensors and limits apoptosis in macrophages. Proc Natl Acad Sci USA 114, E4251– E4260. 10.1073/pnas.1618310114.

38. Klein, S., Cortese, M., Winter, S.L., Wachsmuth-Melm, M., Neufeldt, C.J., Cerikan, B., Stanifer, M.L., Boulant, S., Bartenschlager, R., and Chlanda, P. (2020). SARS-CoV-2 structure and replication characterized by in situ cryo-electron tomography. Nat. Commun. 11, 5885. 10.1038/s41467-020-19619-7.

39. Knoops, K., Kikkert, M., Worm, S.H.E. van den, Zevenhoven-Dobbe, J.C., van der Meer, Y., Koster, A.J., Mommaas, A.M., and Snijder, E.J. (2008). SARS-coronavirus replication is supported by a reticulovesicular network of modified endoplasmic reticulum. PLoS Biol. 6, e226. 10.1371/journal.pbio.0060226.

40. Lemaire, P.A., Anderson, E., Lary, J., and Cole, J.L. (2008). Mechanism of PKR Activation by dsRNA. J. Mol. Biol. 381, 351–360. 10.1016/j.jmb.2008.05.056.

41. Bevilacqua, P.C., and Cech, T.R. (1996). Minor-groove recognition of double-stranded RNA by the double-stranded RNA-binding domain from the RNA-activated protein kinase PKR. Biochemistry 35, 9983–9994. 10.1021/bi9607259.

42. Manche, L., Green, S.R., Schmedt, C., and Mathews, M.B. (1992). Interactions between double-stranded RNA regulators and the protein kinase DAI. Mol. Cell. Biol. 12, 5238– 5248. 10.1128/mcb.12.11.5238-5248.1992.

43. Kindler, E., Gil-Cruz, C., Spanier, J., Li, Y., Wilhelm, J., Rabouw, H.H., Züst, R., Hwang, M., V’kovski, P., Stalder, H., et al. (2017). Early endonuclease-mediated evasion of RNA sensing ensures efficient coronavirus replication. PLoS Pathog. 13, e1006195. 10.1371/journal.ppat.1006195.

44. Burke, J.M., Lester, E.T., Tauber, D., and Parker, R. (2020). RNase L promotes the formation of unique ribonucleoprotein granules distinct from stress granules. J. Biol. Chem. 295, 1426–1438. 10.1074/jbc.RA119.011638.

45. Burke, J.M., Moon, S.L., Matheny, T., and Parker, R. (2019). RNase L Reprograms Translation by Widespread mRNA Turnover Escaped by Antiviral mRNAs. Mol. Cell 75, 1203–1217.e5. 10.1016/j.molcel.2019.07.029.

