## Supplemental Figures for "MERS-CoV antagonizes PKR activation by inhibiting its condensation at viral replication complexes": Supplemental Figures.docx

Contents:

Supplemental figures 1-11

Supplemental videos legends 1-5

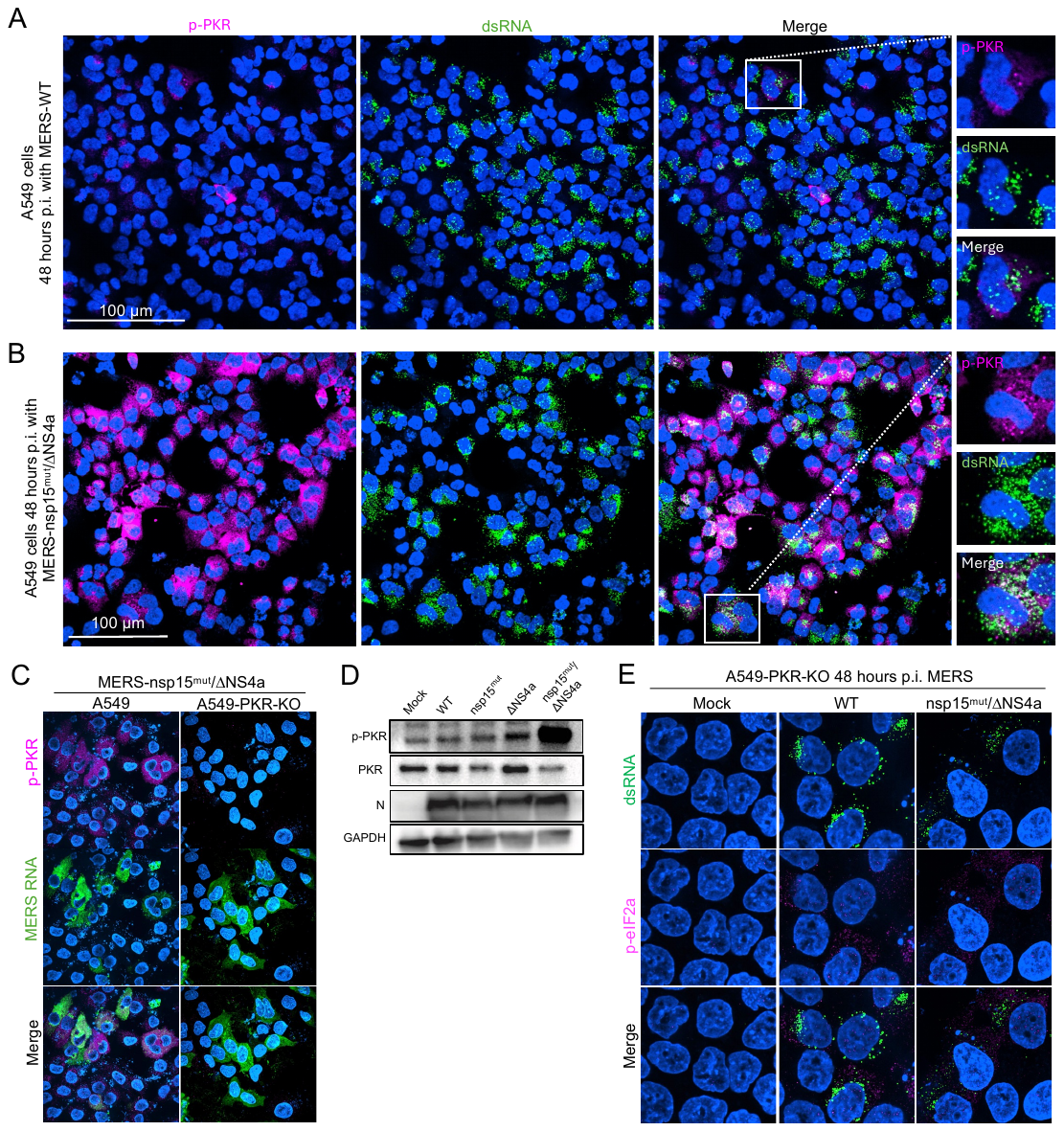
**Figure S1. Increased p-PKR and p-eIF2α levels in MERS-nsp15^mut^/∆NS4a infected cells depend on the presence of PKR.** (A, B) Large field IFA of A549-DPP4 cells infected with (A) MERS-WT (B) MERS-nsp15^mut^/∆NS4a at 48 hours p.i. P-PKR (magenta), and dsRNA (green). (C) IFA combined with smFISH of cells infected with MERS-nsp15^mut^/∆NS4a for 48 hours, with p-PKR (magenta) and 5’ MERS RNA (green). (D) Western blot showing protein levels for indicated proteins at 48 hpi. (E) IFA for p-eIF2α (magenta) and dsRNA (green) in PKR-KO cells 48 hpi.

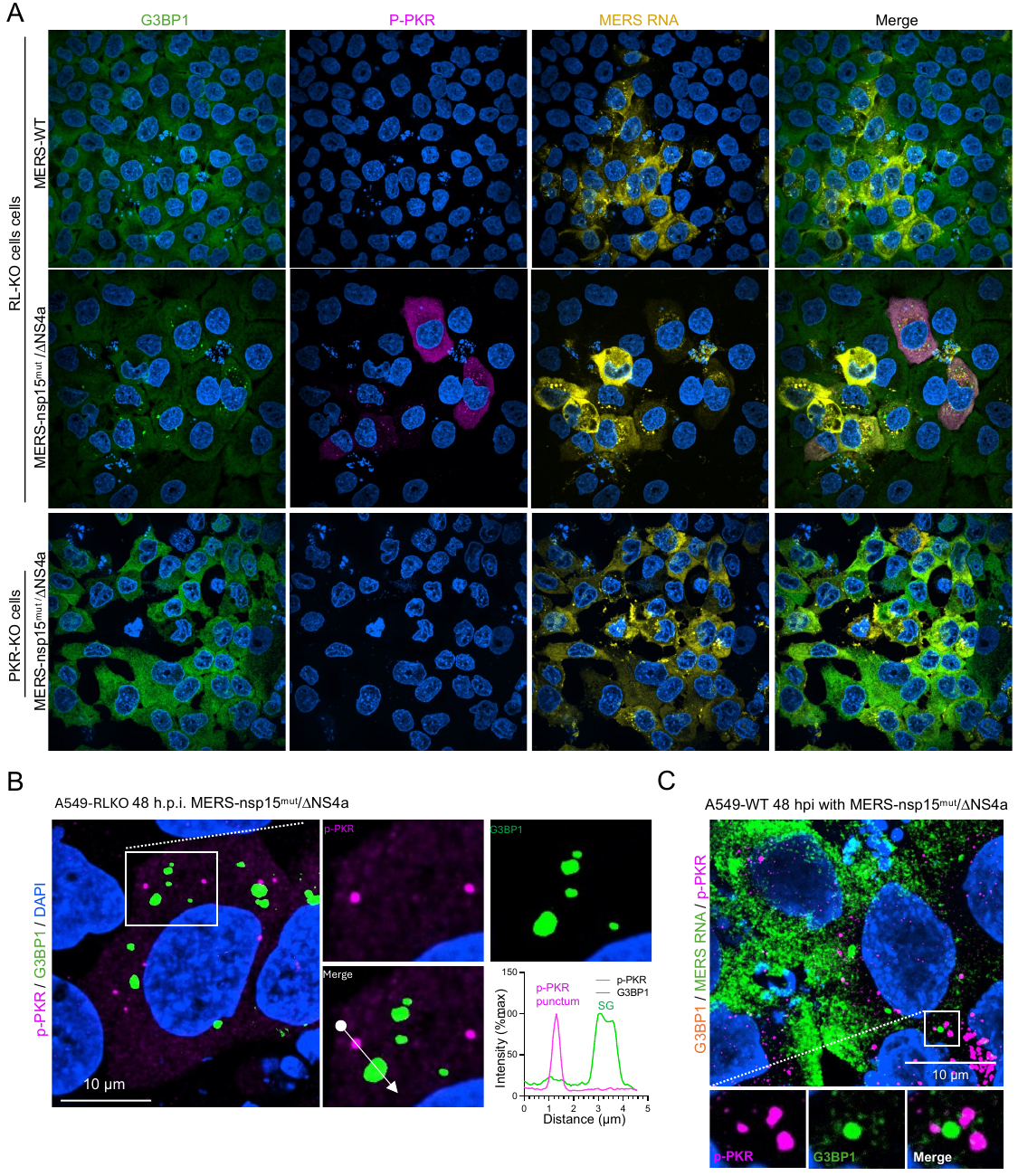
**Figure S2. PKR puncta correlate to stress granule assembly but are distinct from G3BP1 ribonucleoprotein complexes, including stress granules and RNase L induced-bodies.** (A) IFA combined with smFISH of RL-KO and PKR-KO A549-DPP4 cells infected with MERS-WT or MERS-nsp15^mut^/∆NS4a for 48 hours. G3BP1 is shown in green, p-PKR in magenta, and MERS RNA is shown in yellow. (B) IFA on A549-DPP4-RL-KO cells infected with MERS-nsp15^mut^/∆NS4a for 48h and stained for p-PKR (magenta) and G3BP1 (green). Line trace depicts IFA signal through the line shown in the merged inset. (C) IFA combined with smFISH of A549-DPP4 cells infected with MERS-nsp15^mut^/∆NS4a for 48h. Nuclei were stained with DAPI (blue) for all images.

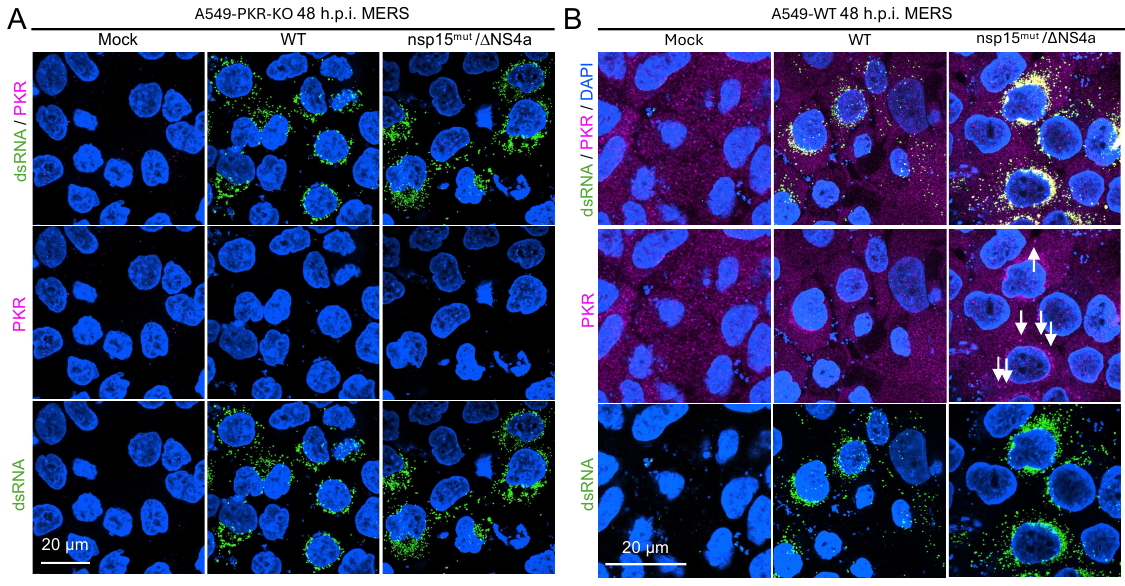

**Figure S3. The PKR puncta are PKR-dependent and observed with total PKR antibody.** (A) IFA showing PKR (magenta) and dsRNA (green) in A549-DPP4-PKR-KO cells 48 hpi. (B) IFA showing PKR (magenta) and dsRNA (green) in A549-DPP4 cells 48 hpi. Arrows indicate PKR puncta.

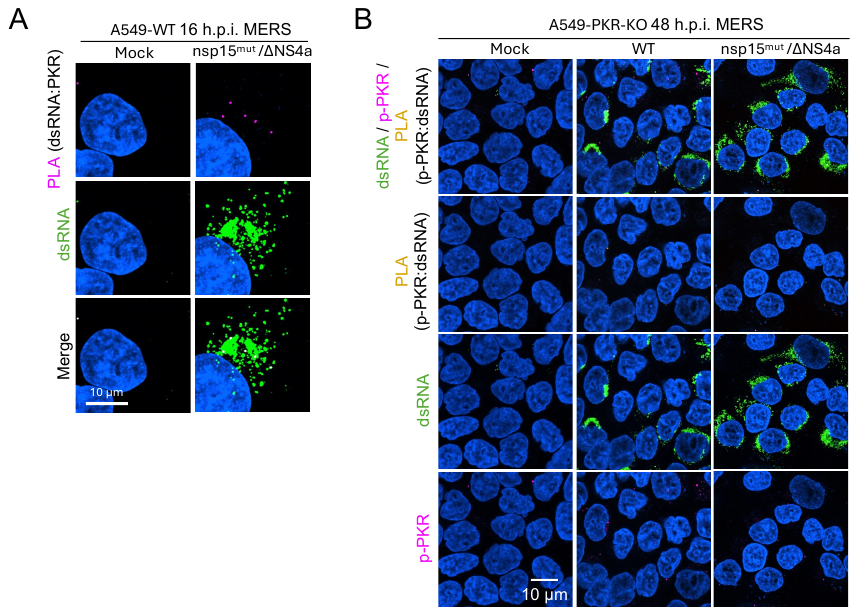

**Figure S4. Proximity-ligation assay for dsRNA and PKR.** (A) PLA (PKR : dsRNA) (magenta) in A549-DPP4 cells either mock treated or infected with indicated MERS-nsp15^mut^ /∆NS4a virus for 16 hours. Co-IFA was done for dsRNA (green). (B) PLA (p-PKR : dsRNA) (yellow) conducted in A549-DPP4-PKR-KO cells either mock treated or infected with indicated MERS virus for 48 hours. Co-IFA was done for dsRNA (green) and p-PKR (magenta).

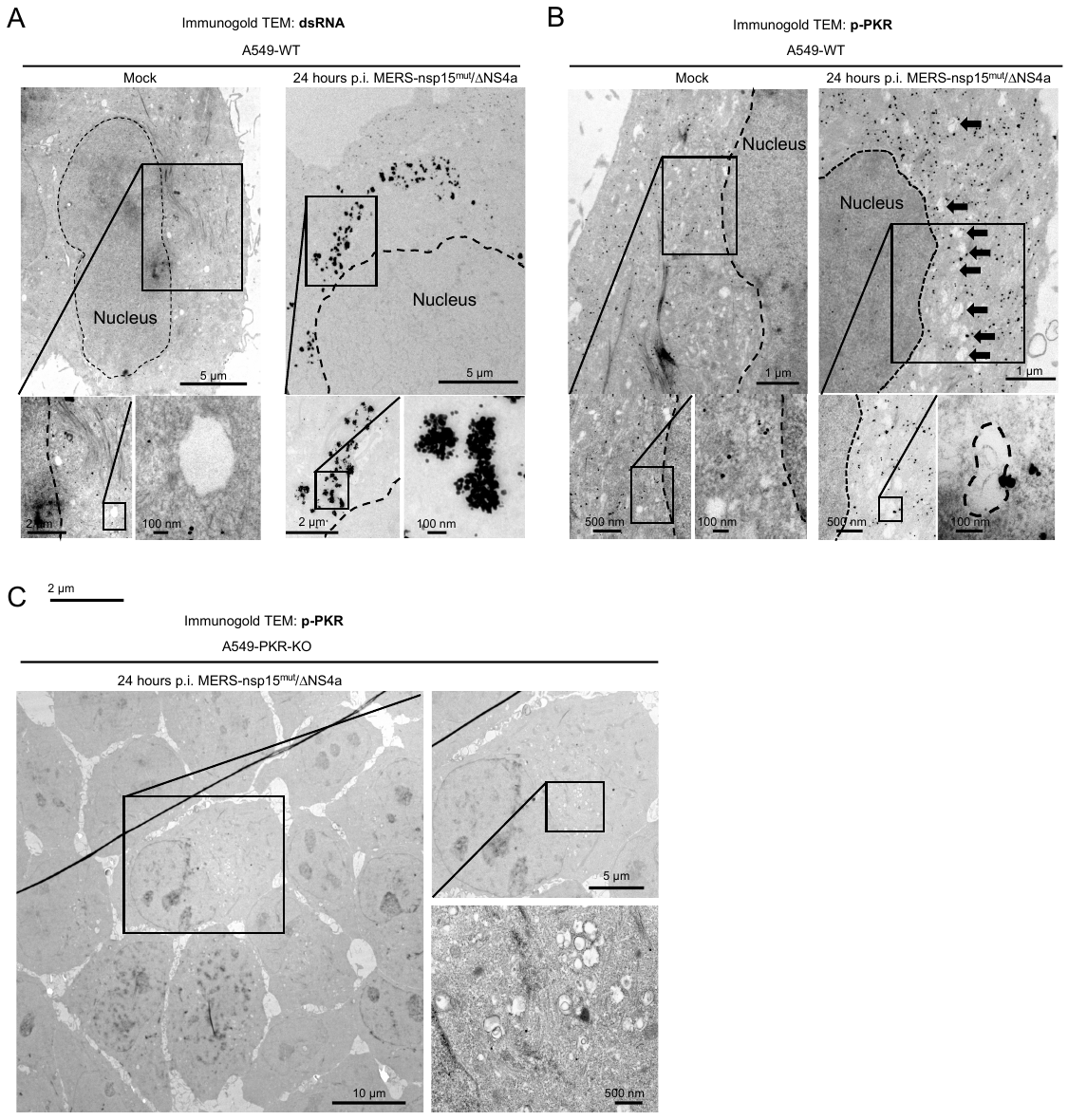

**Figure S5. Mock cells lack viral DMVs and immunogold staining for dsRNA, and p-PKR signal is dependent on expression of PKR.** (A-C) Transmission electron microscopy of A549-DPP4 cells 24 hours post infection. (A) Immunogold staining for dsRNA in WT cells either mock treated or 24 hours p.i. with MERS-nsp15^mut^/∆NS4a. (B) Immunogold staining for p-PKR in WT cells either mock treated or 24 hours p.i. with MERS-nsp15^mut^/∆NS4a. Arrows indicate DMVs with p-PKR condensates associated to the structure. (C) Immunogold staining for p-PKR in PKR-KO cells either mock treated or 24 hours p.i. with MERS-nsp15^mut^/∆NS4a.

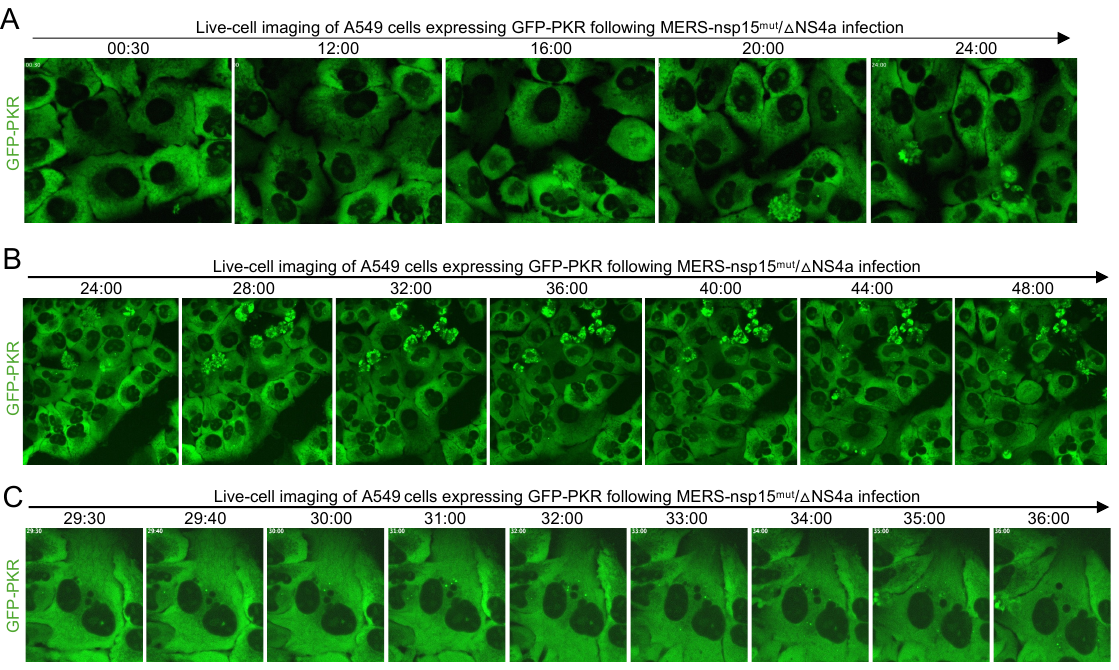

**Figure S6. Live cell imaging of MERS-nsp15mut/∆NS4a infected cells show the dynamics of PKR-DRIFs.** (A-C) Live cell imaging of A549-DPP4 cells expressing GFP-tagged PKR during the course of MERS-nsp15^mut^/∆NS4a infection. (A) Cells imaged from 0.5 hours to 24 hours p.i. (B) Cells imaged from 24 hours to 48 hours p.i. (C) Cells imaged at 29 hours 30 minutes until 36 hours.

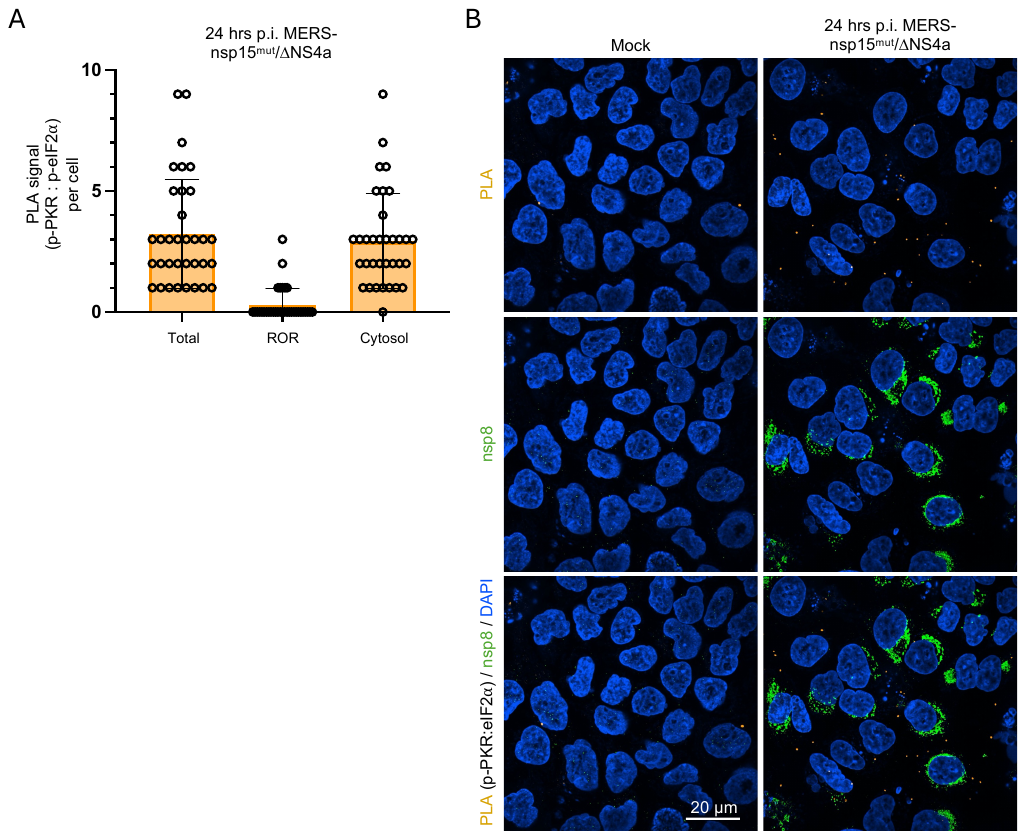

**Figure S7. PLA signal between eIF2α and p-PKR is primarily cytosolic.** (A) Expanded quantification of Figure 4D, showing the localization of PLA signal in cells infected with MERS-nsp15^mut^/∆NS4a for 24 hours. (B) Large field images of PLA (eIF2α : p-PKR) signal (yellow) in A549-DPP4 cells at 24 hours p.i. with MERS-nsp15^mut^/∆NS4a. Co-IFA signal for nsp8 (green).

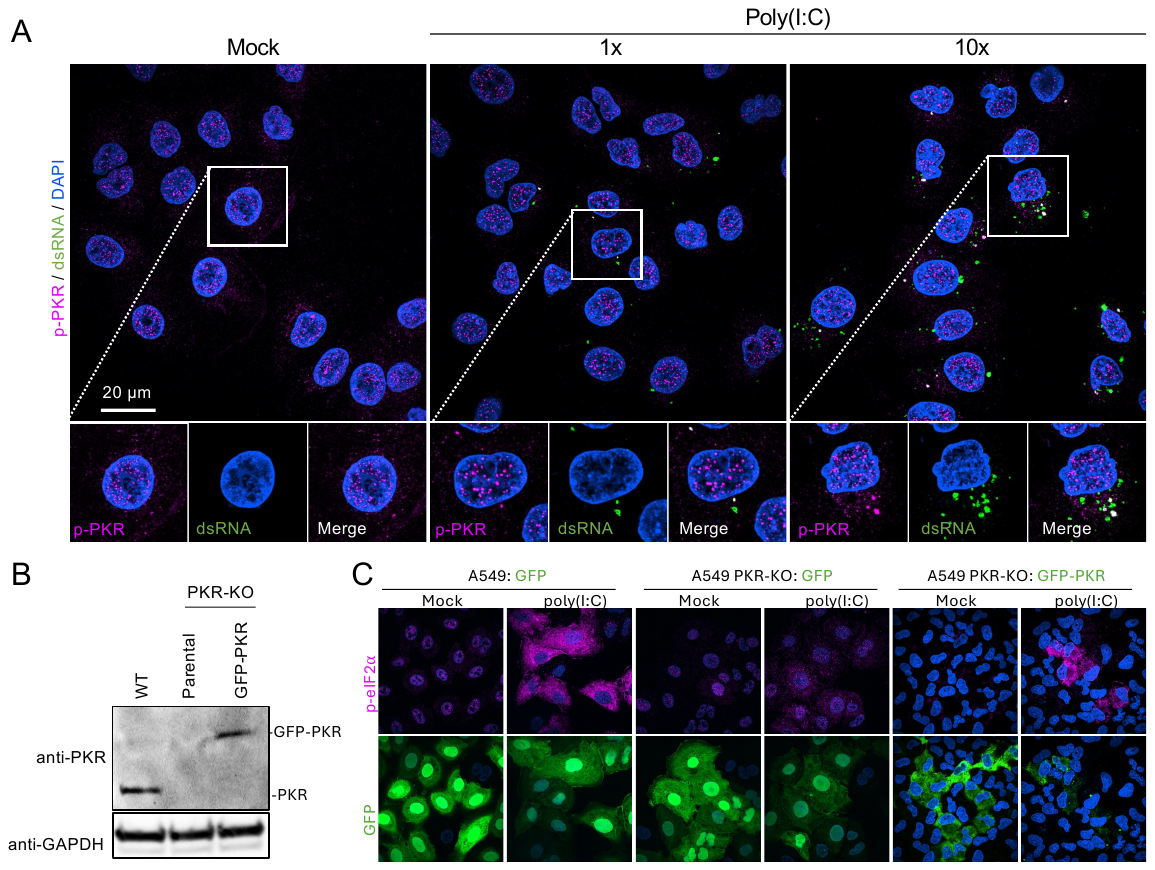

**Figure S8. Increasing amounts of dsRNA results in increased number of PKR-DRIFs. (**A) IFA showing p-PKR (magenta) and dsRNA (green) in A549 cells treated with Poly(I:C) for 6 hours at 1x (250ng) or 10x (2.5 mg) concentration. (B) Western blot of parental A549 cells or A549-PKR-KO cell with or without GFP-PKR expression.

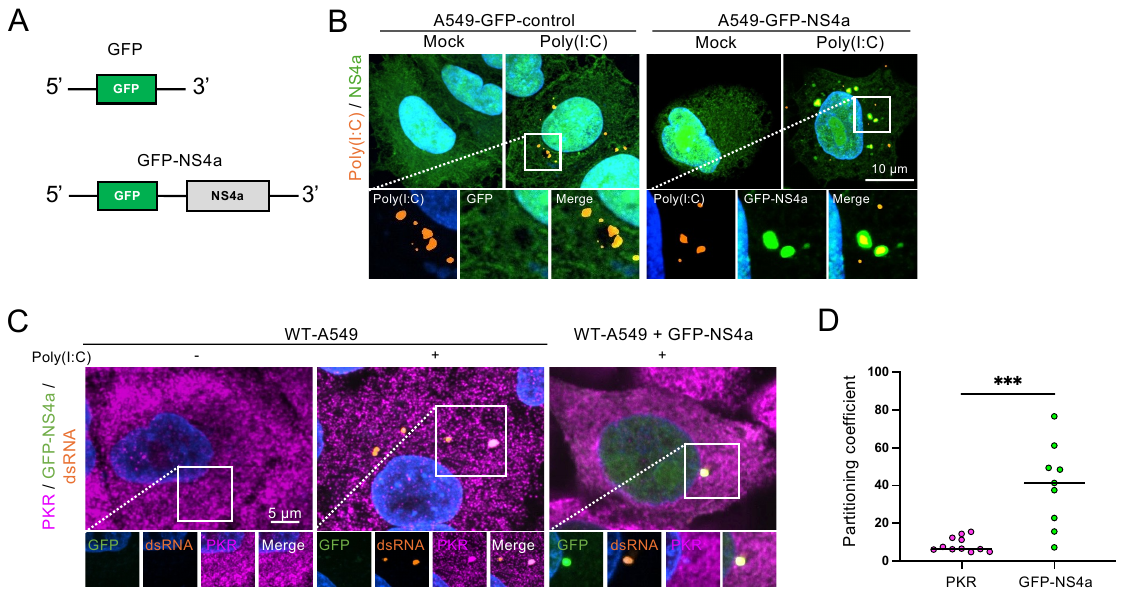

**Figure S9. A549 cell lines expressing GFP-control and GFP-NS4a.** (A) Design of A549 cell lines expressing either GFP-control and GFP-NS4a. (B) IFA showing p-PKR (magenta) and Rhodamine-Poly(I:C) (orange) in A549 cells, or cell lines expressing GFP-control or GFP-NS4a. (C) IFA showing PKR (magenta) and Rhodamine-Poly(I:C) (orange) in A549-WT with or without expressing GFP-NS4a, treated with or without Poly(:C) for 6 hours. (D) Quantification for the partitioning coefficient from (C), where each dot is one condensate.

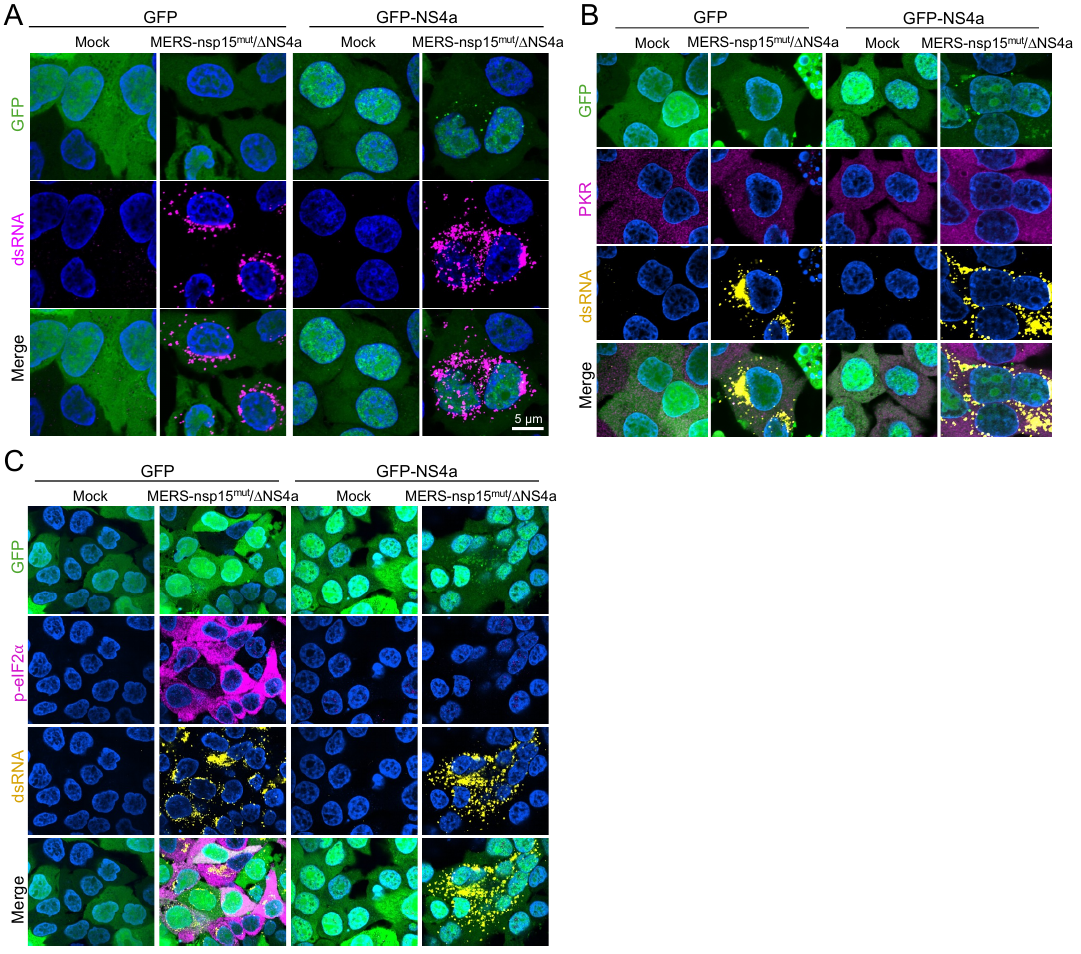

**Figure S10. NS4a inhibits PKR-DRIF assembly and PKR activation.** (A) IFA for dsRNA in A549-DPP4 cells expressing either GFP or GFP-NS4a following 24 hrs. p.i. MERS-nsp15^mut^/∆NS4a or mock treatment. (B) IFA for dsRNA and PKR in A549-DPP4 cells expressing either GFP or GFP-NS4a following 24 hrs. p.i. MERS-nsp15^mut^/∆NS4a or mock treatment. (C) IFA for dsRNA and p-eIF2⍺ in A549 cells expressing either GFP or GFP-NS4a following 24 hrs. p.i. MERS-nsp15^mut^/∆NS4a or mock treatment.

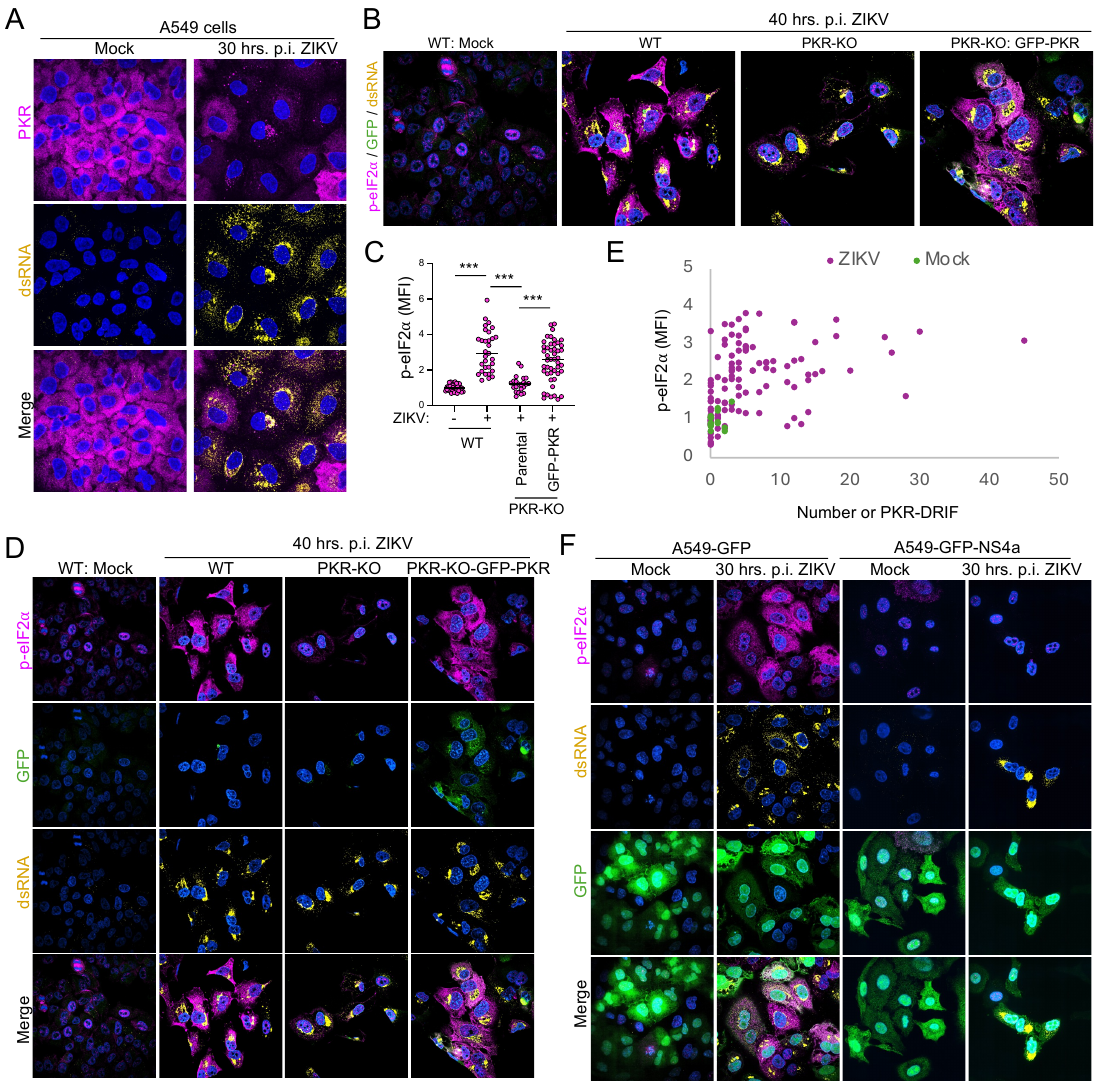

**Figure S11. PKR-DRIF assembly correlates to PKR activation in response to ZIKV infection.** (A) IFA for dsRNA(yellow) and PKR (magenta) in A549 cells 30 hrs. p.i. ZIKV or mock treatment. (B) IFA for dsRNA (yellow) and p-eIF2⍺ (magenta) in either WT or PKR-KO A549 cells with or without expressing GFP-PKR following 40 hrs. p.i. ZIKV or mock treatment. (C) Quantification of the mean fluorescence intensity (MFI) of p-eIF2α per cell (dot) as represented in (B and D). (D) Large field images of IFA for dsRNA (yellow) and p-eIF2⍺ (magenta) in either WT or PKR-KO A549 cells expressing GFP-PKR following 40 hrs. p.i. ZIKV or mock treatment. (E) Correlation plot showing p-eIF2⍺ levels and number of PKR-DRIF. (F) IFA of A549 cells expressing either GFP-control or GFP-NS4a mock treated or post 30-hour ZIKV infection. Cells were stained for p-eIF2⍺ (magenta) and dsRNA (yellow).

**Supplemental Video 1 (related to Fig S6A).**

Live cell imaging of A549-DPP4 cells expressing GFP-PKR 0-24 hrs. p.i. MERS-nsp15^mut^/∆NS4a. Images were taken every 10 min. starting at the time of infection. Time p.i. is indicated on upper left corner of the panel.

**Supplemental Video 2 (related to Fig S6B).**

Live cell imaging of A549-DPP4 cells expressing GFP-PKR 24-48 hrs. p.i. MERS-nsp15^mut^/∆NS4a. Images were taken every 10 min. starting at 24 hrs. p.i. Time p.i. is indicated on upper left corner of the panel.

**Supplemental Video 3 (related to Fig 3E).**

Live cell imaging of A549-DPP4 cells expressing GFP-PKR 24-48 hrs. p.i. MERS-nsp15^mut^/∆NS4a. Images were taken every 10 min. starting at 24 hrs. p.i. Time p.i. is indicated on upper left corner of the panel.

**Supplemental Video 4 (related to Fig 7D).**

Live cell imaging of A549-PKR-KO cells expressing GFP-PKR 24-48 hrs. p.i. ZIKV. Time p.i. is indicated on upper left corner of the panel.

**Supplemental Video 5 (related to Fig 7D).**

Live cell imaging of A549-PKR-KO cells expressing GFP-PKR 24-48 hrs. p.i. ZIKV. Images were taken every 10 min. starting at 24 hrs. p.i. Time p.i. is indicated on upper left corner of the panel.
